# Sliding windows analysis can undo the effects of preprocessing when applied to fMRI data

**DOI:** 10.1101/2023.10.06.561221

**Authors:** Martin A. Lindquist

## Abstract

Resting-state fMRI (rs-fMRI) data is used to study the intrinsic functional connectivity (FC) in the human brain. In the past decade, interest has focused on studying the temporal dynamics of FC on short timescales, ranging from seconds to minutes. These studies of time-varying FC (TVFC) have enabled the classification of whole-brain dynamic FC profiles into distinct “brain states”, defined as recurring whole-brain connectivity profiles reliably observed across subjects and sessions. The analysis of rs-fMRI data is complicated by the fact that the measured BOLD signal consists of changes induced by neuronal activation, as well as non-neuronal nuisance fluctuations that should be removed prior to further analysis. Thus, the data undergoes significant preprocesing prior to analysis. In previous work [24], we illustrated the potential pitfalls involved with using modular preprocessing pipelines, showing how later preprocessing steps can *reintroduce* correlation with signal previously removed from the data. Here we show that the problem runs deeper, and that certain statistical analysis techniques can potentially interact with preprocessing and reintroduce correlations with previously removed signal. One such technique is the popular sliding window analysis, used to compute TVFC. In this paper, we discuss the problem both theoretically and empirically in application to test-retest rs-fMRI data. Importantly, we show that we are able to obtain essentially the same brain states and state transitions when analyzing motion induced signal as we do when analyzing the preprocessed but windowed data. Our results cast doubt on whether the estimated brain states obtained using sliding window analysis are neuronal in nature, or simply reflect non-neuronal nuisance signal variation (e.g., motion).

## 1 Introduction

Resting-state fMRI (rs-fMRI) is used to study the intrinsic functional connectivity (FC) in the human brain ([5]). It has been used to show that fluctuations in the blood oxygen level dependent (BOLD) signal in spatially distant brain regions are strongly and consistently correlated with one another ([3, 14]). Originally, statistical methods used to access FC assumed it was constant across a given experimental run (i.e., between 5-15 minutes long). However, in recent years, interest has extended to studying the temporal dynamics of FC on much shorter timescales (i.e., seconds to minutes) ([20, 29, 26]). These studies of time-varying FC (TVFC), also known as dynamic connectivity, have enabled the classification of whole-brain dynamic FC profiles into distinct “brain states”, defined as recurring whole-brain connectivity profiles reliably observed across subjects throughout the course of a resting state run ([9]).

The analysis of rs-fMRI data is complicated by the fact that the measured BOLD signal consists of changes induced by neuronal activation (the signal of interest) and non-neuronal variation. Examples of the latter include drift, spiking artifacts, motion-related artifacts, and fluctuations due to physiological sources (e.g., heart rate and respiration) ([28, 35, 4, 33]). Failure to properly control and correct for these types of variation can have significant impact on subsequent analysis, as they can potentially induce spurious FC between different brain regions. Thus, it is of great interest to remove or at least reduce their effects on the analysis of rs-fMRI data.

For these reasons, rs-fMRI data are subjected to a series of preprocessing steps ([8, 11, 38]) prior to statistical analysis. Many preprocessing pipelines used in the field are modular in nature, by which we mean they are composed of a number of separate distinct filtering/regression steps. These include, among other steps, removal of head motion covariates using linear regression and band-pass filtering, performed sequentially and in a flexible order. In previous work ([24]), we illustrated the potential pitfalls involved with using modular preprocessing pipelines for data cleaning. We showed how later preprocessing steps can *reintroduce* correlation with signal removed from the data in prior preprocessing steps. In particular, we illustrated how performing high-pass filtering after motion regression can bring motion induced variation back into the data. Similarly, performing motion regression after high-pass filtering can bring unwanted frequency components back into the signal; a finding previously pointed out by [18].

However, it is important to stress that it is not only preprocessing steps that can interact with one another in such a manner, but perhaps more insidiously statistical analysis techniques can similarly interact with previous preprocessing steps and reintroduce correlation with previously removed signal. For example, we hypothesize that a statistical analysis technique that incorporates temporal filtering, that either removes or attenuates certain frequencies in the time series data, may also bring back motion induced signal variation in an analogous manner as outlined above.

One such technique commonly used in the analysis of rs-fMRI data is the so-called sliding windows analysis ([1, 10, 19]), used to determine TVFC. Here a series of correlation matrices are computed over fixed-length, windowed segments of the fMRI time series. The relationship between sliding window correlation-based TVFC estimates and nuisance factors was previously explored in [27]. There it was found that TVFC was correlated with temporal fluctuations in the norm of the nuisance regressors, and that this correlation remained even after nuisance regression was performed. We hypothesize that this is in part be due to interactions between the application of the window and nuisance regression. Further, they showed that block regression where nuisance regression is performed separately within each window is similarly ineffective in removing nuisance effects from TVFC estimates. We hypothesize that this is in part be due to the manner in which nuisance regression effects higher-order statistics.

In this paper we investigate these issues in detail. We begin by outlining the theoretical concerns, and show how it is possible to reintroduce correlation to signal previously removed from the data during preprocessing. We thereafter illustrate the problem empirically using test-retest rs-fMRI data. We utilize a standard processing pipeline used to determine brain states in a TVFC study, and show that the derived brain states are corrupted by motion induced signal variation previously removed during preprocessing. Importantly, we show that we are able to obtain essentially the same brain states and transitions when analyzing motion-related signal as we do when analyzing the preprocessed but windowed rs-fMRI data. These results cast doubt on whether the estimated brain states obtained using sliding windows analysis are neuronal in nature, or simply reflect non-neuronal nuisance signal variation. Finally, we present some partial fixes to the problem.

## 2 Methods

### 2.1 Computing time-varying functional connectivity

Typically, TVFC between multiple regions of the brain are represented using a series of either covariance or correlation matrices, that depict the relationship between different brain regions or components over time. The most common approach for estimating the elements of these matrices is to use the sliding windows approach. Here, a time window of fixed length *L* is selected, and data points within that window are used to calculate the metric of interest (e.g., correlation coefficient). The window is then shifted across time, using overlapping windows, and a new value for the metric is computed at each time point. Throughout we assume that the window is shifted by one TR at a time.

To illustrate, consider the sliding-window covariance between two time series **x** and **y**. The repeated calculation of the local covariance obtained by shifting a window *w* across time can be written:

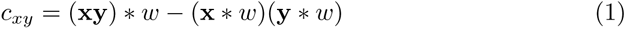

where ∗ denotes the convolution operator. Using the Fourier convolution theorem, this can be expressed in the frequency domain as:

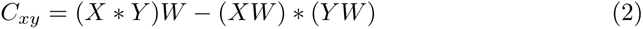

where capital letters represent the DFT of the respective signals.

If *w* were sinc-shaped, convolution with a sliding window in the time-domain is equivalent to applying a low-pass filter in the frequency domain and would have a similar effect. We therefore hypothesize that this operation will negatively interact with signal removed in previous preprocessing steps in an analogous manner outlined in [24]. However, in practice one does not use a sinc-shaped window when computing time-varying connectivity. Instead, a rectangular window of length *L* is commonly used. The DFT of a rectangular window is given by the discrete version of a sinc-function, whose main lobe has width 1*/L*. Hence, in this setting it is no longer a low-pass filter, but rather a filter that attenuates high frequency components of the signal. This attenuation is increased if a tapered window is used instead of a rectangular window.

In this paper we primarily consider sliding-window correlations, which are normalized functions of the covariance. The normalization is determined by the reciprocal of the square root of the product of (**x**^2^) ∗ *w* and (**y**^2^) ∗ *w* as follows:

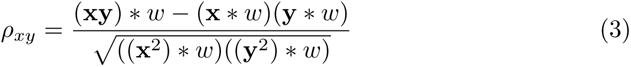

TVFC can be computed using data from individual voxels ([20, 23]), averaged over pre-specified regions of interest ([10]), or estimated using data-driven methods such as independent component analysis ([1]). Once the set of time-varying correlations have been estimated, a standard approach for summarizing them is to classify them into distinct brain states, or recurring whole-brain patterns of FC that appear repeatedly across time and subjects ([1]). To derive brain states, the time-varying correlations across subjects are clustered, typically using k-means ([1]). Practically, this is done by reorganizing the lower triangular portion of each subject’s *d* × *d* × *T* time-varying correlation data into a matrix with dimensions ((*d* − 1)*d/*2) × *T*, where *d* is the number of voxels/regions/components and *T* the number of time points. The data from all subjects is then concatenated into a matrix with dimensions ((*d* − 1)*d/*2) × *TN*, where *N* is the number of subjects. K-means clustering is performed and each of the resulting cluster centroids represent a recurring brain state. The number of clusters can be chosen by computing the within-group sum of squares for each candidate number of clusters and picking the ‘elbow’ in the plot; though alternative approaches exist.

### 2.2 Understanding the interaction between sliding window correlation and nuisance regression

As seen in Eq. 3, the time-varying correlation between two time series **x** and **y** depends on a number of different components. These include the windowed time series (**x** ∗ *w* and **y** ∗ *w*), their product ((**x** ∗ *w*)(**y** ∗ *w*)), the windowed Hadamard product ((**xy**) ∗ *w*), and the windowed (non-normalized) standard deviations ((**x**^2^)∗*w* and (**y**^2^)∗*w*). In the following section we study the effects of windowing on both a single time series, as well as the product of two time series. This will allow us to investigate the effects of windowing on each component of the time-varying correlation.

We begin with studying the effects on a single time series.

Let **y** be an *n*-dimensional vector consisting of rs-fMRI signal from a single voxel in the brain. Further, let **X** be an *n* × *p* design matrix whose columns consist of the nuisance regressors we seek to remove from **y**. This can be done by fitting the following linear model:

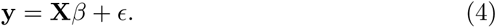

The least-squares estimate of *β* is given by

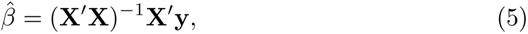

and the fitted value by:

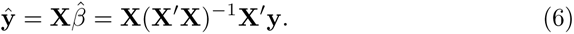

Here **ŷ** represents the estimated nuisance signal in the voxel and corresponds to signal that we seek to remove from **y**.

To simplify calculations, we take a projection approach towards linear regression. We define a projection matrix

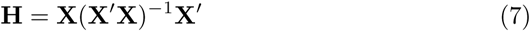

and write **ŷ** = **Hy**. Here **H** is a linear matrix transformation that projects **y** onto the space spanned by the columns of the design matrix **X**. An important property of projection matrices are that they are both idempotent (**H** = **H**^2^) and symmetric (**H** = **H***^′^*).

To remove the effects of motion we subtract **ŷ** from the data and compute the residual vector **e** = **y** − **ŷ**. This can alternatively be expressed **e** = (**I** − **H**)**y**, and this is our new signal of interest. Note **I** − **H** is also a projection matrix which projects the data onto a subspace orthogonal to the columns of the design matrix **X**. Thus, our signal of interest **e** lies in a subspace orthogonal to **X**. Hence,

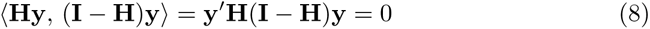

which implies that the data is uncorrelated with the nuisance signal. Here ⟨*., .*⟩ represents the Euclidean inner product, and the last equality holds since **H** = **H**^2^. At this point we have, to the extent possible using linear projections, removed the effects of the nuisance components in **X** from the data.

Next, suppose we apply a sliding window to the data.

This can be represented as the application of a Toeplitz matrix operator **W** to the signal, written **We**. Unfortunately, this operation may move the data into a subspace no longer orthogonal to the space spanned by **X**. This would have the effect of partially reintroducing a relationship between the nuisance signal and the signal being studied. This can be seen by observing that

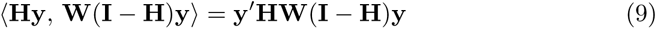

is no longer guaranteed to be zero. In other words, because the operator **W** is not restricted to operate in the sub-space spanned by **I** − **H**, Eq. (9) may not be zero. This is illustrated in Fig. 1. Similarly, there may be a relationship between the windowed nuisance signal and the signal being studied. Note in the continuation we use the expressions for the convolution **x** ∗ *w* and Toeplitz matrix operation **Wx** interchangeably.

**Figure 1:**
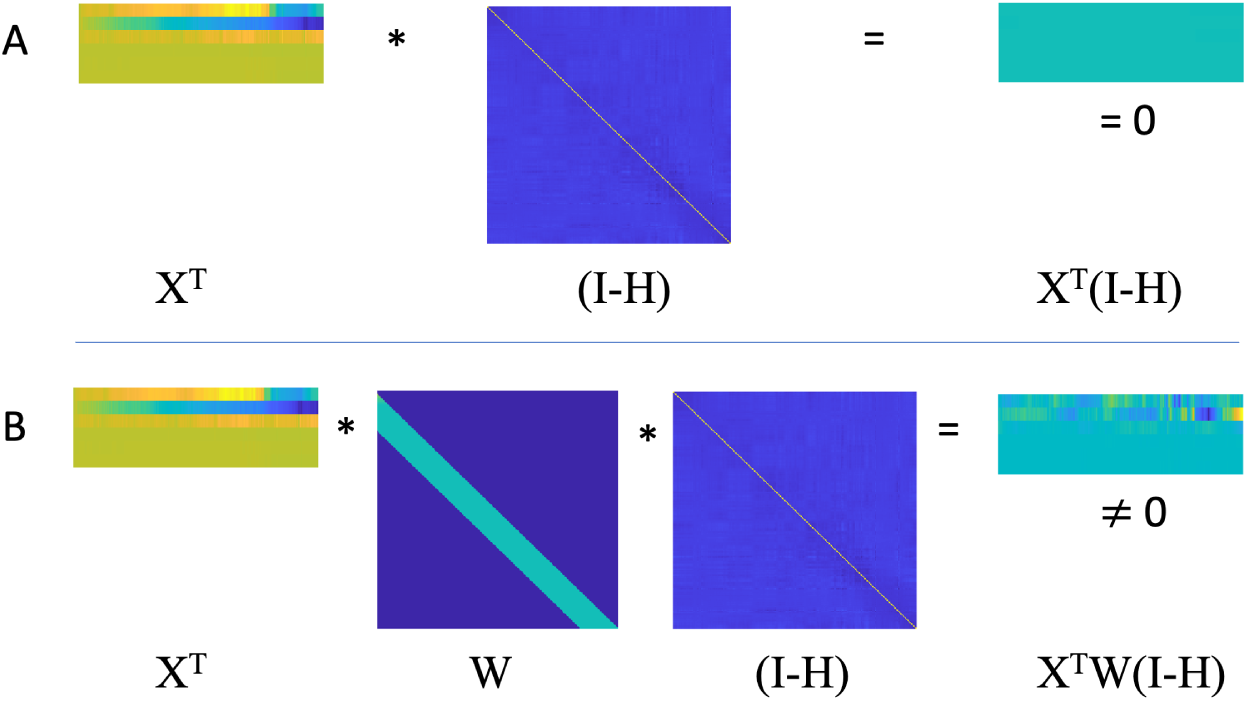
A graphical illustration of the effect of multiple matrix operations. Let **X** be a design matrix whose columns consist of the nuisance regressors and **H** = **I** − **X**(**X***^′^***X**)*^−^*^1^**X***^′^* be a projection matrix. Further, let **W** be a Toeplitz matrix that applies a sliding window. In this setting, the following two results hold: (A) The columns of **X** are orthogonal to **I** − **H** as shown by the fact that the product **X***^′^*(**I** − **H**) = **0**; (B) The columns of **X** are not guaranteed to be orthogonal to **W**(**I**−**H**) as shown by the fact that the product **X***^′^***W**(**I**−**H**) ≠ **0**.

Fig. 2A illustrates this relationship geometrically. Consider a vector **y** (the measured signal) and a vector **z** (the nuisance signal). After projecting **y** onto **z**, we obtain the vector **e**, which is the nuisance-corrected signal. As anticipated, it is orthogonal to **z**. This is equivalent to the results seen in Eq. 8. However, the application of a Toeplitz matrix **W**, rotates and scales **We**. The resulting vector is no longer required to be orthogonal to **e**. This is equivalent to the results seen in Eq. 9. Hence, the application of **W** has the effect of potentially reintroducing correlation with nuisance-induced signal variation into the data.

**Figure 2:**
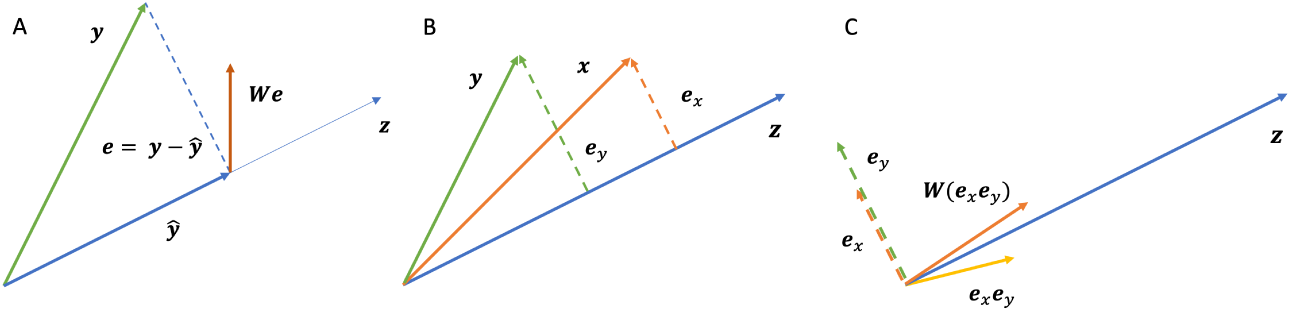
A geometric illustration of the effect of multiple matrix operations. (A) The vector **y** in 2-dimensional space (green line) is projected onto the vector **z**, which gives the vectors **ỹ** (blue line) and **e** = **y**−**ŷ** (dashed line). Importantly, **e** is orthogonal to **z**. After applying the matrix **W**, the resulting vector **We** (red line) is no orthogonal to **z**. (B) Consider two vectors **x** and **y** (orange and green lines, respectively). Using the same procedure outlined in (A), the vectors **e***_x_* and **e***_y_* are both orthogonal to **z**. (C) The Hadamard product **e***_x_***e***_y_* (yellow line) is no longer orthogonal to **z**, nor is **W**(**e***_x_***e***_y_*) (red line).

The description above relates to the effects of windowing on the terms **x** ∗ *w* and **y** ∗ *w* in Eq. 3. However, it is also important to investigate the effects on the product of two time series, particularly the term (**xy**) ∗ *w*. In Fig. 2B, suppose we expand the example to include two vectors **x** and **y**. After projecting both vectors on **z**, we obtain the vectors **e_x_** and **e_y_**, respectively, which are the nuisance-corrected signal obtained from **x** and **y**. As anticipated by theory, both are orthogonal to **z**. In panel C, consider taking the Hadamard product **e_x_e_y_**. This vector is no longer orthogonal to **z**. In addition, the application of a Toeplitz matrix **W**, complicates the issue further by rotating and scaling the Hadamard product. This operation also alters the correlation with **z**. Using the same reasoning, one can see the same results will hold when considering the terms (**x**^2^) ∗ *w* and (**y**^2^) ∗ *w*.

In sum, the application of a sliding-window can reintroduce a correlation between nuisance signal and the terms **x** ∗ *w*, **y** ∗ *w*, (**xy**) ∗ *w*, (**x**^2^) ∗ *w* and (**y**^2^) ∗ *w* in Eq. 3. In addition, correlation can also be reintroduced with (**xy**) ∗ *w* (and other products of time series) due to the fact that nuisance regression is not designed to be orthogonal to higher-order statistics. Hence, we hypothesize that the correlation between nuisance signal and time-varying connectivity depends on a complex relationship between windowing and the effects of taking the Hadamard product between the two nuisance-corrected signals.

### 2.3 Data collection

We used the Multi-Modal MRI Reproducibility Resource (Kirby) from the F.M. Kirby Research Center^1^; see [21] for more information about the acquisition protocol. Briefly, it consists of data from 20 healthy participants scanned on a 3T Philips Achieva scanner. Each participant completed two scanning sessions on the same day, and between sessions a full repositioning of the participant occurred. A T1-weighted MPRAGE structural image was acquired during both sessions (acquisition time = 6 *min*, TR/TE/TI = 6.7*/*3.1*/*842 *ms*, resolution = 1 × 1 × 1.2 *mm*^3^, SENSE factor = 2, flip angle = 8*^◦^*). A multi-slice SENSE-EPI pulse sequence ([34, 30]) was used to acquire a single rs-fMRI run during each session, consisting of 210 volumes sampled every 2 *s* at 3 *mm* isotropic spatial resolution (acquisition time: 7 *min*, TE = 30 *ms*, SENSE acceleration factor = 2, flip angle = 75*^◦^*, 37 axial slices collected sequentially with a 1 *mm* gap). Participants were instructed to rest comfortably while remaining as still as possible, and no other instruction was provided. We refer to the first rs-fMRI session collected as Session 1 and the second as Session 2.

### 2.4 Preprocessing

SPM 12 (Wellcome Trust Centre for Neuroimaging, London, United Kingdom) and custom MATLAB (The Mathworks, Inc., Natick, MA) was used to preprocess the data. To allow for the stabilization of magnetization, four volumes were discarded at acquisition, and an additional volume was discarded prior to preprocessing. Slice timing correction was performed using the slice acquired at the middle of the TR as reference, and rigid body realignment parameters were estimated to adjust for head motion. The six rigid-body parameters were saved for further analysis; see below. Structural runs were registered to the first functional frame and spatially normalized to Montreal Neurological Institute (MNI) space using SPM’s unified segmentation-normalization algorithm ([2]). The estimated rigid body and nonlinear spatial transformations were applied to the rs-fMRI data.

### 2.5 Creation of nuisance images and nusiance norm

Before further preprocessing, we estimated the total amount of motion at each voxel of the brain for each subject and session. To do so we created a design matrix **X** consisting of 24 regressors. These included the six motion regressors obtained after rigid-body transformation, the regressors squared, their first order difference, and the first order difference squared. Together these 24 regressors have been used to correct for so-called spin-history artifacts (i.e., residual effects of motion after rigid-body correction) ([17]). The fitted values of this motion regression are typically removed from the signal prior to further analysis, giving rise to what we will refer to as motion-corrected images. Fig. 3 provides a graphical illustration of the approach.

**Figure 3:**
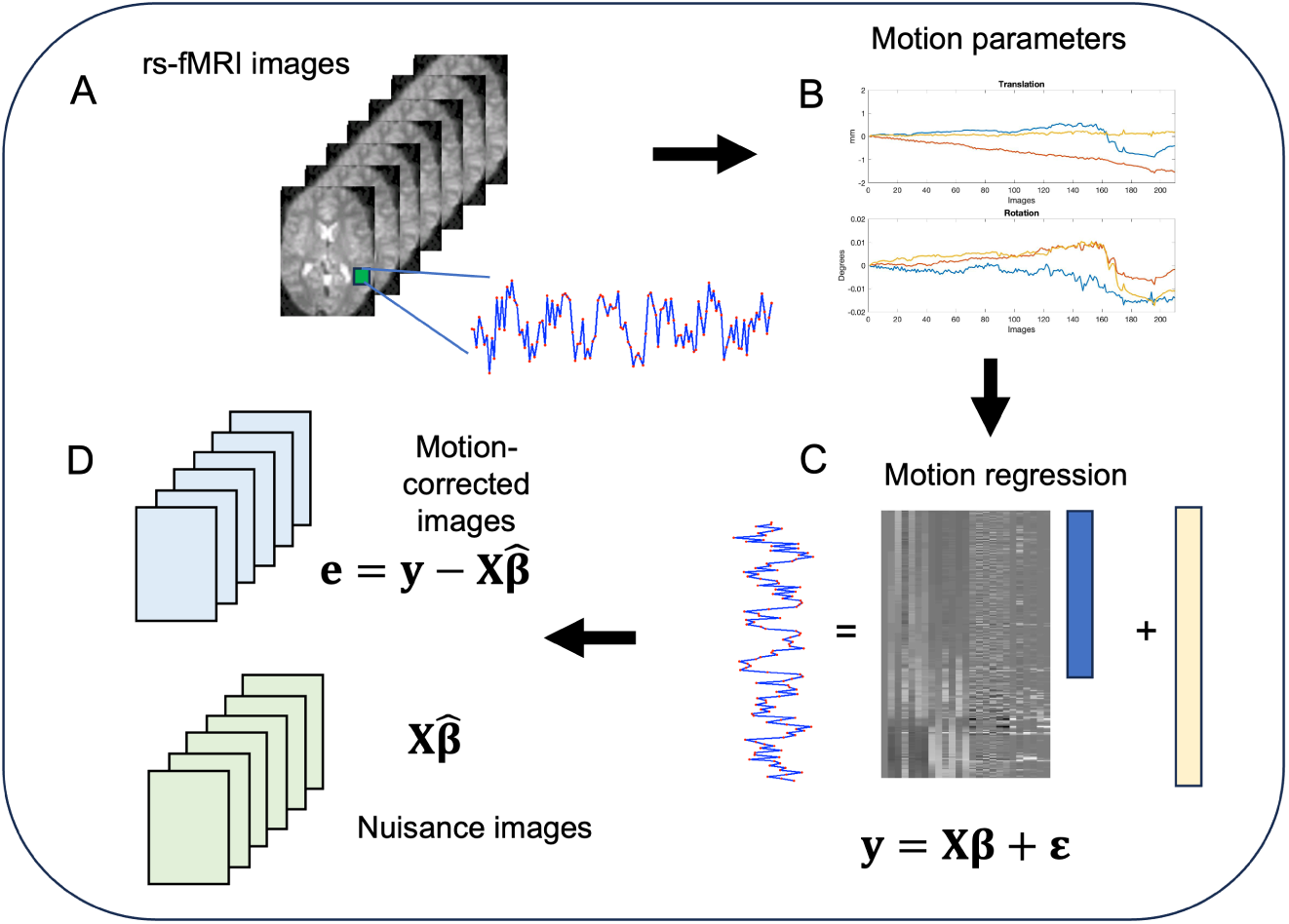
An illustration of the procedure used to create nuisance images. (A) A signal is extracted from a single voxel over time. (B) Prior to extraction of the signal a rigid body realignment is performed to adjust for head motion, and the six parameters were saved for further analysis. (C) The six motion parameters obtained after rigid-body transformation, the parameters squared, their first order difference, and the first order difference squared are used as predictors in a linear model to remove additional motion-related artifacts in a process known as motion regression. (D) In the motion-corrected images, the effects of motion are removed from the signal of interest and used to create new data orthogonal to this confounder. In the nuisance images the fitted values from the motion regression are used to create new data consisting purely of nuisance signal.

Here **y** corresponds to the signal of interest from voxel *v*, and **X** is the design matrix containing the 24 motion regressors. To remove the effect of motion we fit the motion regression **y** = **X***β* + *ɛ*. After estimating the parameters *β* corresponding to the motion regressors at each voxel *v*, we use them to create ‘nuisance images’. This was done by computing the fitted values **X***β* at each voxel, thereby creating 4D images of the estimated motion induced signal variation at each voxel in the brain; see Fig. 3D. These nuisance images can be extended to incorporate other non-neuronal signal (e.g., signal from white matter or cerebral spinal fluid), but we only consider motion here for simplicity. These images provide a summary of the information related to nuisance covariates removed from the signal.

To investigate the relationship between the nuisance signal and TVFC estimates, following [27] we used the windowed norm of the nuisance regressors. It is computed by first demeaning each of the *i* = 1*, … N_R_* nuisance regressors in each window *k*. Let (*n_i_*(*t_k_*)*, … , n_i_*(*t_k_* + *L* − 1)), be the *i^th^* demeaned nuisance time course in the *k^th^* window where *t_k_* is the starting time of the window and *L* the window length. The nuisance norm for the *k^th^* window is given by:

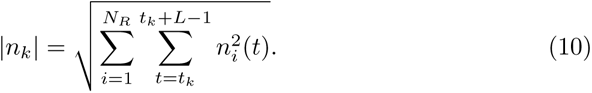

The norm is computed for all windows by temporally shifting the window index *k*, giving rise to a nuisance norm time course. To examine the relationship between the nuisance norm and TVFC estimates, we computed the correlation coefficient between the two time-varying correlations.

The nuisance norm is a subject-specific measure. To obtain a connection-specific measure of the relationship between the nuisance signal and TVFC estimates, we also examined the relationship between the TVFC estimates and the time-varying correlation between the motion components extracted from **x** and **y**, which we denote *e***_x_** and *e***_y_**, respectively. As above, this is performed by computing the correlation coefficient between the two time-varying correlations.

### 2.6 Analysis pipelines

Next we applied six different analytic pipelines to the data, outlined below.

#### Pipeline 1

In the first pipeline, we parcellated the data into 268 non-overlapping regions using the Shen atlas ([32]), and performed sliding windows correlation and k-means clustering. We used a window length of 30 TRs, which is the suggested optimal window-length for rs-fMRI data collected using a standard EPI sequence with a sampling frequency of 2 *s* ([23]). K-means clustering was repeated 50 times, using random initialization of centroid positions, to increase the chance of escaping local minima. The elbow method was used to choose the two cluster solution, corresponding to two reoccurring brain states. This was repeated for both sessions for each of the 20 subjects. Since we did not perform motion regression in this pipeline the effects of motion should still be present in the signal on which TVFC was assessed.

#### Pipeline 2

In the second pipeline, we repeated the same procedure described in Pipeline 1, but first performed motion regression (as described in Section 2.5) and removed the estimated motion from the rs-fMRI data prior to parcellating the data.

#### Pipeline 3

In the third pipeline, we repeated the same procedure described in Pipeline 1 on the nuisance images. Importantly, these images consist entirely of motion induced signal variation that was removed in Pipeline 1. We use this pipeline to get a clearer picture of the effects of motion on the estimated TVFC.

#### Pipeline 4

In the fourth pipeline we used the same data as in Pipeline 1, but performed a secondary motion regression procedure inside of each sliding window. This approach, referred to as block regression [27], provides a way to mitigate the effects of motion artifacts when using a sliding window. We use this approach as it allows motion regression and application of the sliding window to be performed simultaneously, so not to interact with one another. Similar to [27] we used the six standard motion parameters (three translations, three rotations) as our regressors.

#### Pipeline 5

In the fifth pipeline we used the same data as in Pipeline 1, but applied a ‘motion-corrected window’ rather than a standard rectangular window. This window is defined as

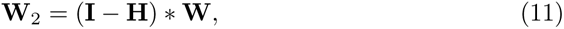

and was created to ensure that the windowing matrix is orthogonal to the space spanned by the motion regressors. Here

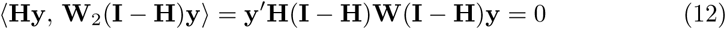

which implies that the data is uncorrelated with the motion signal.

#### Pipeline 6

The sixth pipeline was equivalent to Pipeline 4, with the difference that twelve motion parameters were used instead of six. This included the six standard motion parameters and the regressors squared to help model quadratic motion related effects.

### 2.7 Assessing the Impact of Motion

To investigated the impact of motion at each step of the analysis we compute the average fitted value from the motion regression over each region (corresponding to the regional averages of the nuisance images). We computed the correlation between this signal and the average rs-fMRI over each region both with and without motion regression. The former corresponds to Pipeline 2 and the latter to Pipeline 1. We further computed the correlation between the motion signal and the resting-state signal after application of a sliding window kernel to show that it reintroduces a correlation with previously removed motion induced signal variation. Finally, we repeated the procedure using the block regression procedure with both six and twelve regressors (Pipelines 4 and 6), and using the motion corrected window defined in Eq. 11.

Next, we calculated the correlation between the nuisance norm and the components of time-varying connectivity for each subject and pair-wise connection. This included the correlation between nuisance norm and the time-varying values of (**xy**) ∗ *w*, (**x** ∗ *w*)(**y** ∗ *w*), **x** ∗ *w* and (**x**^2^) ∗ *w*, where **x** and **y** are the signals of interest. This procedure was repeated using the time-varying correlation between the motion extracted from the time series pair **x** and **y**, which can be obtained from Pipeline 3, instead of the nuisance norm.

After applying k-means clustering to the TVFC estimates obtained from each pipeline, individual time points were allocated to either one of the two reoccurring brain states (clusters). We used custom Matlab code to determine transition times between states for each participant. This allowed for the estimation of the number of transitions and dwell time within a given state for each participant. The distribution for the number of transitions was modeled using a Poison distribution whose parameter *λ* was estimated from the data. This distribution is often used to model the number of events (in this case transitions) occurring within a given interval of time. The distribution for the dwell time was modeled using an exponential distribution with parameter *µ*. This distribution is often used to model the time between events in a Poisson process.

Next, the brain states estimated using the six different pipelines were compared using multidimensional scaling (MDS). MDS provides a distance-preserving map of the input data in two-dimensional space, thus providing a visual representation of the distance between brain states. This allows us to evaluate ‘how close’ brain states obtained using the various pipelines lie to one another.

Finally, to assess the impact of motion on Pipeline 2, which is the standard pipeline, we estimated the proportion of motion-induced signal variation after windowing using the following metric: 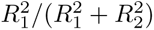. Here 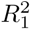 is the squared correlation between the motion data and the windowed time series expressed as *corr*(**Hy**, **P**(**I** − **H**)**y**) and 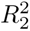 is the squared correlation between the motion-corrected data and the windowed time series expressed as *corr*((**I** − **H**)**y**, **P**(**I** − **H**)**y**). This metric allowed us to investigate the proportion of motion induced signal variation present in the signal after the windowing procedure. Prior to windowing, this metric will be 0 in all regions of the brain.

### 2.8 Simulation

We performed a simple simulation study to investigate the relationship between the components of the estimated time-varying correlation and motion. We defined the true time-varying correlation at time *t* as *ρ*(*t*) = .5 ∗ *sin*(2 ∗ *π* ∗ *t/*150). Hence, the values oscillate between −0.5 and 0.5 in a periodic manner; see Fig. 4A. Next, for each of the 20 subjects included in the study, we computed the subject-specific nuisance norm time courses (see Fig. 4B) and a subject-specific motion-corrected windowing matrix according to Eq. 11.

**Figure 4:**
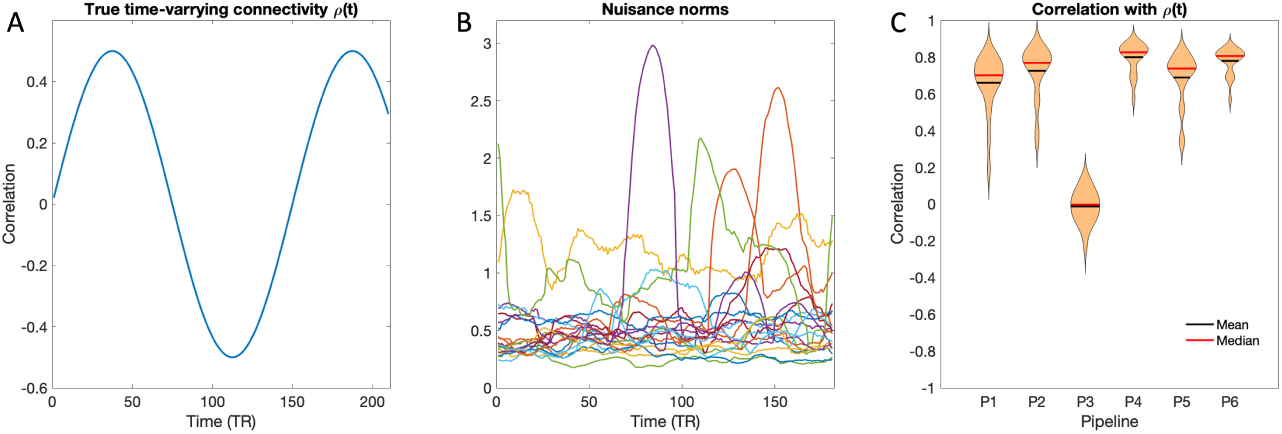
Simulation study. (A) The true time-varying correlation *ρ*(*t*) at time *t*. (B) The subject-specific nuisance norm time courses for each of the 20 subjects. (C) Violin plots of the mean correlation between the estimated time-varying connectivity and the true time-varying connectivity *ρ* plotted across the 20 subjects for each of the six pipelines.

For each ‘subject’ we completed 1000 repetitions of the simulation. In each, we generated two time series from a bivariate normal distribution with mean 0, variance 1, and correlation *ρ*(*t*). Next, we extracted two motion time series, randomly chosen from two of the 268 possible regions. These motion time series (estimated using 6 regressors) were combined with the simulated time series to create two time series **x** and **y**, designed to include both a true time-varying correlation *ρ*(*t*) and a realistic motion component.

For each repletion, we estimated the time-varying connectivity for the six different pipelines and computed its correlation with *ρ*(*t*) to assess pipeline performance. In addition, we computed the correlation between the subject-specific nuisance norms and the time-varying correlation and covariance values between **x** and **y**. Finally, we computed the correlation between the nuisance norm and the windowed values of **x**, **y**, **xy**, **x**^2^, and **y**^2^. We repeated this process using the time-varying correlation between the motion components of **x** and **y** (i.e., the signal used in Pipeline 3), in place of the nuisance norm.

Fig. 4C shows violin plots of the mean correlation (across repetitions) between the estimated time-varying connectivity and the true time-varying connectivity *ρ* plotted across the 20 subjects for each of six pipelines. Here, the pipelines using block regression (Pipelines 4 and 6) perform best, and the difference between them is negligible. The motion data (Pipeline 3) has the worst performance with a mean correlation centered around 0.

Fig. 5A shows the correlation between the windowed time series and the nuisance norm plotted across the 1000 repetitions for an example subject. Interestingly, the correlation between the nuisance norm and the time-varying connectivity are roughly similar to those observed between the nuisance norm and the time-varying covariance and the windowed *xy*-product. In contrast, the correlation between the nuisance norm and **x**, **y**, **x**^2^, and **y**^2^, are all roughly centered around 0, with the latter two showing a slight positive bias. Together, these findings indicate that the correlation between the time-varying correlation and the nuisance norm is largely driven by the **xy** product. These results are consistent across other subjects (not shown here). Fig. 5B shows equivalent violin plots comparing the mean correlations over the 1000 repetitions plotted across all 20 subjects. Again, there are clear similarities between the correlation, covariance, and **xy** product. The (**x**) ∗ *w* and (**y**) ∗ *w* terms are consistently centered around zero for all subjects, as indicated by their small variance. However, the (**x**^2^) ∗ *w* and (**y**^2^) ∗ *w* terms show a positive bias that somewhat shifts the distribution of the the correlation compared to the covariance.

**Figure 5:**
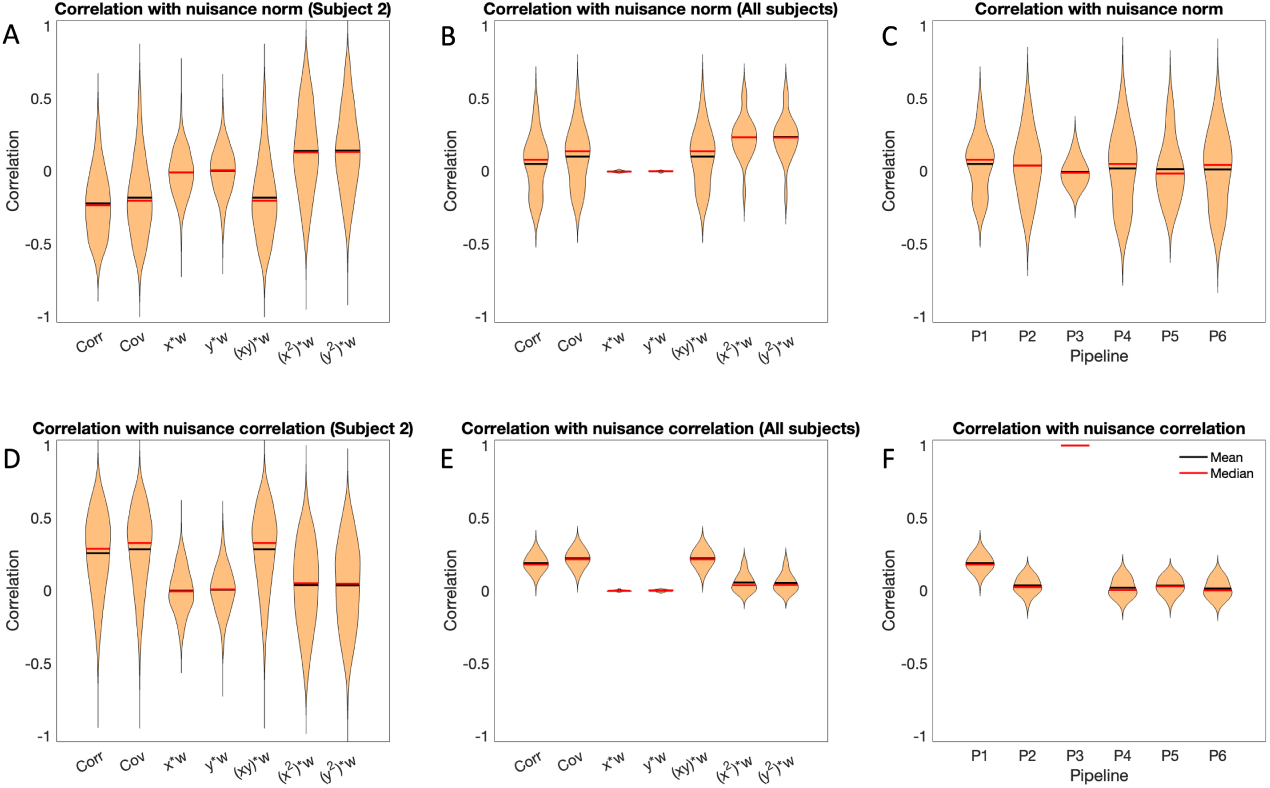
Results of the simulation study. (A) Violin plots of the correlation between windowed time series and the nuisance norm plotted across the 1000 repetitions for an example subject. (B) Violin plots of the mean correlation between windowed time series and the nuisance norm over the 1000 repetitions plotted across the 20 subjects. (C) Violin plots of the mean correlation between the estimated time-varying connectivity and the nuisance norm over the 1000 repetitions plotted across the 20 subjects for each of the six pipelines. (D-F) Equivalent results using the time-varying correlation between motion time series (i.e., the results from Pipeline 3) instead of the nuisance norm.

Finally, Fig. 5C shows violin plots of the mean correlation between the estimated time-varying connectivity and the nuisance norm plotted across the 20 subjects for each of the six pipelines. Surprisingly, the pure motion data (Pipeline 3) has the lowest correlation with the nuisance norm. This is presumably due to the fact that the relationship between motion and the motion norm is not linear. It does raise the point that the correlation with the nuisance norm, while important, may not tell the full story about the relationship between TVFC and motion.

For this reason, we also compute the correlation with the time-varying correlation between the extracted motion signals, i.e. the results obtained from Pipeline 3. These correlations are shown in Fig. 5D-F. Interestingly, the bias in the (**x**^2^) ∗ *w* and (**y**^2^) ∗ *w* terms disappear; see Panels D-E. Studying Panel F, the mean correlation obtained using Pipeline 3 is by definition equal to 1. In addition, apart from Pipeline 1, the remaining four perform equivalently.

## 3 Results

We begin with an empirical illustration of the effects of windowing on a single time series. Fig. 6A shows the correlation between the signal and motion in each region using the data where no motion regression was performed (Pipeline 1). Clearly, the signal is highly correlated with motion, showing that motion will play a prominent role in the signal without additional preprocessing. Panel B shows the same correlation after motion regression was performed (Pipeline 2). In this setting, as expected all motion has been properly removed from the signal. However, Panel C shows the correlation between the windowed motion corrected time course and the motion. Here the windowing process has led to a reintroduction of motion induced signal variation and motion effects are once more correlated with the signal of interest. Finally, Panel D shows the results between the windowed time course and the motion, in the setting where motion regression is performed within each window. This procedure has led to a significant reduction in correlation, though not a complete removal; see Section 4 for further discussion.

**Figure 6:**
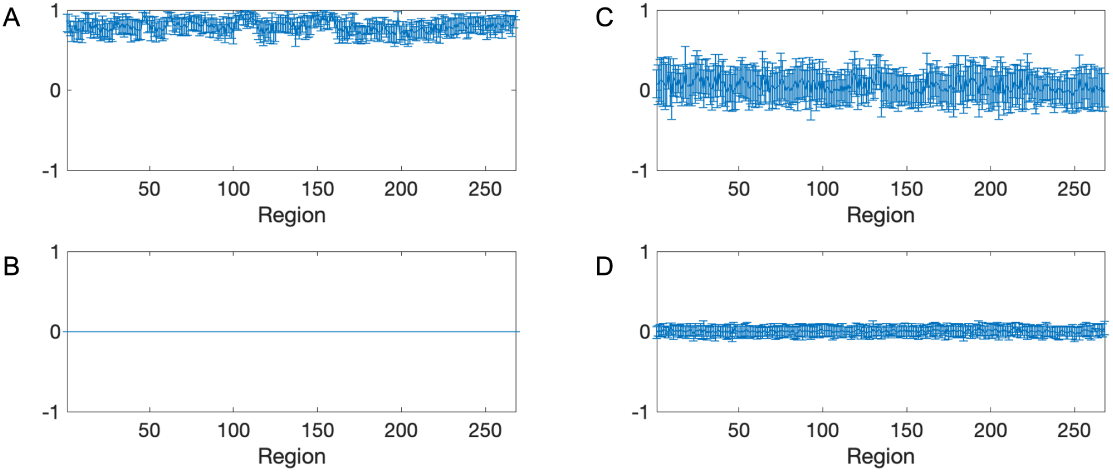
Empirical illustration. (A) The correlation between the motion signal and the non-motion corrected rs-fMRI signal over each region (Pipeline 1). (B) The correlation between the motion signal and the motion-corrected rs-fMRI signal over each region (Pipeline 2). (C) The correlation between the motion signal and the resting-state signal after the application of a sliding window kernel (Pipeline 2); illustrating the reintroduction of motion induced signal variation after applying the sliding window. (D) The correlation between the motion signal and the resting-state signal after the application of a sliding window kernel and secondary motion regression (Pipeline 4). In all figures the vertical bars represent one standard error.

Fig. 7 shows three different Manhattan-style plots of the correlation between motion and windowed rs-fMRI signal for Session 1. Panel A shows the correlation between the motion signal and the windowed rs-fMRI signal over each region for each subject and pipeline. Panel B shows the correlation between the windowed time series and the windowed motion. Finally, Panel C shows how the windowed time course is correlated with the nuisance norm.

**Figure 7:**
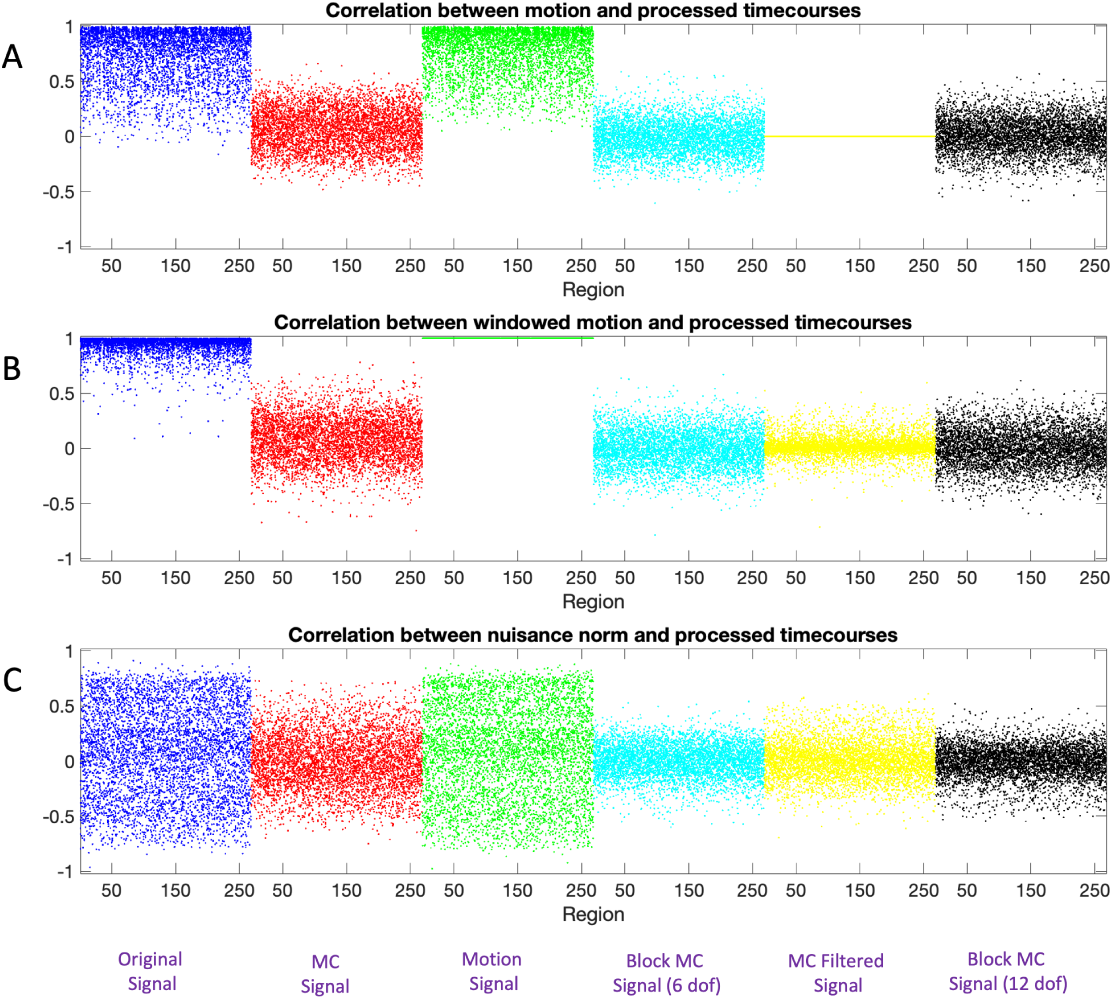
Correlation between motion and windowed rs-fMRI signal for Session 1. (A) The correlation between the motion signal and the windowed rs-fMRI signal for each of the 20 subjects shown for each region and pipeline. Note there are 20 dots representing the subject-specific values at each point on the x-axis. (B) The correlation between the windowed motion signal and the windowed rs-fMRI signal for each of the 20 subjects shown for each region and pipeline. (C) The correlation between the nuisance norm and the windowed rs-fMRI signal for each of the 20 subjects shown for each region and pipeline. The color-coded as follows: Pipeline 1 (Blue), Pipeline 2 (Red), Pipeline 3 (Green), Pipeline 4 (Cyan), and Pipeline 5 (Yellow).

The results in blue depict the windowed non-motion corrected signals correlation with motion (Pipeline 1). The correlation with the motion time series is generally quite high and the equivalent correlations with the nuisance norm has a large variance. The results in red depict the windowed motion-corrected signals correlation with motion (Pipeline 2). The correlation with the motion time series is substantially lower compared to the non-motion corrected signal. The results in green depict the windowed motion signals correlation with motion (Pipeline 3). Naturally, the windowed motion is perfectly correlated with itself (Panel B), but also highly correlated with the raw motion time course and the nuisance norm. Interestingly, the results are not noticeably different from those seen for the non-motion corrected signal, illustrating how much motion contributes to the raw signal. The results in cyan depict the the correlation between motion and the windowed signals motion-corrected using block regression with 6 regressors (Pipeline 4). The correlations are somewhat less variable than the equivalent results obtained using the motion-corrected signal (Pipeline 2). The results in yellow depict the correlation between motion and the windowed signals obtained using the motion-corrected window function (Pipeline 5). The correlation is zero with the motion time course as it is designed to be orthogonal to the motion. However, the signal is correlated with the windowed motion and nuisance norm at roughly the level seen in Pipeline 4. Finally, the results in cyan depict the correlation between motion and the windowed signals motion-corrected using block regression with 12 regressors (Pipeline 6). The results are roughly equivalent to those seen for Pipeline 4.

These results only represent one part of the calculation of the time-varying connectivity, whose values also depend on the terms (**xy**)∗*w*, (**x**∗*w*)(**y**∗*w*), (**xx**)∗ *w*, and (**yy**) ∗ *w*; see Eq. 3. Fig. 8 shows violin plots of the correlation between the nuisance norm and the various components of time-varying connectivity plotted across all subjects and connections/regions for Session 1.

**Figure 8:**
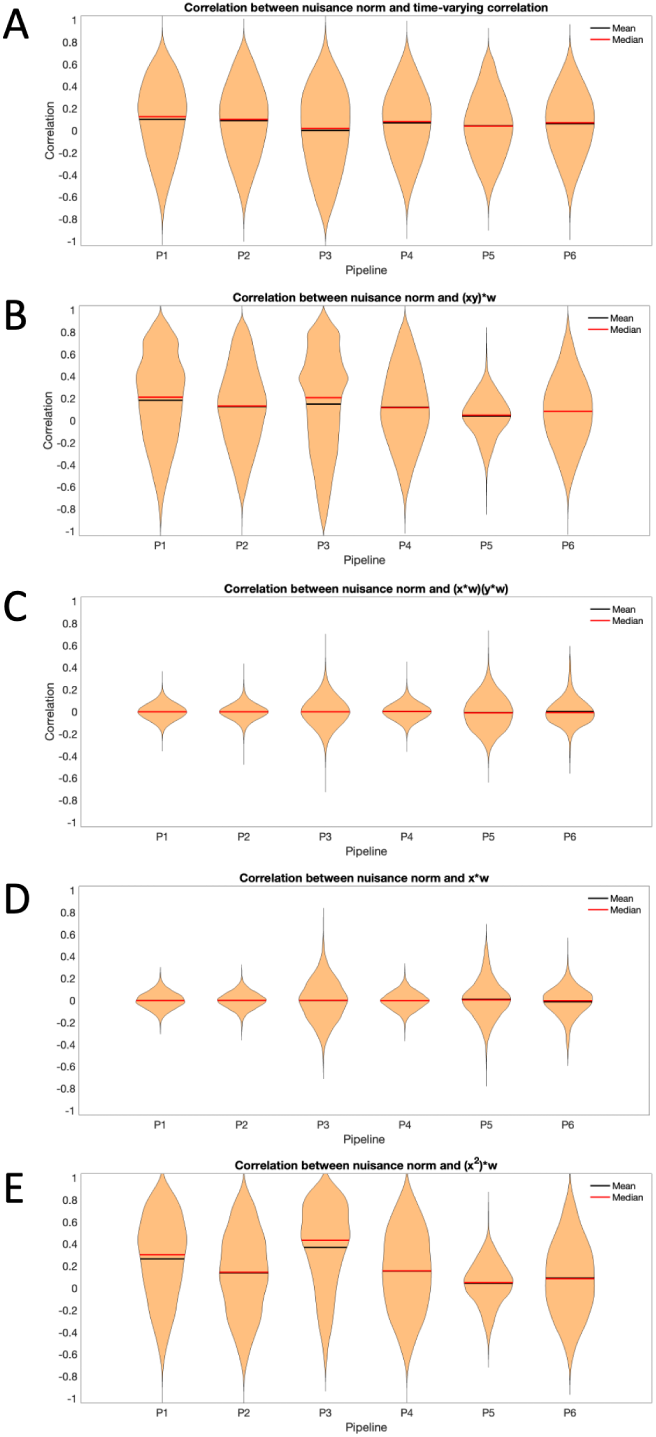
Correlation between the nuisance norm and the components of time-varying connectivity for Session 1. (A) The correlation between the nuisance norm and the time-varying connectivity plotted across all subjects and pair-wise connections. (B) The correlation between the nuisance norm and the time-varying value of (**xy**) ∗ *w*, where **x** and **y** are the signals of interest, plotted across all subjects and pair-wise connections. (C) The correlation between the nuisance norm and the time-varying value of (**x** ∗ *w*)(**y** ∗ *w*) plotted across all subjects and pair-wise connections. (D) The correlation between the nuisance norm and the time-varying value of (**x** ∗ *w*) plotted across all subjects and regions. (E) The correlation between the nuisance norm and the time-varying value of (**x**^2^) ∗ *w* plotted across all subjects and regions.

Panel A shows a violin plot of the correlation between the nuisance norm and the time-varying connectivity plotted across all subjects and pair-wise connections. Clearly there is significant variance for all pipelines, showing the difficulties in completely addressing this issue. Panel B begins to dig into what drives this variation, studying the correlation between nuisance norm and the time-varying value of (**xy**) ∗ *w* plotted across all subjects and pair-wise connections. Clearly most of the variation in the time-varying connectivity is driven by this component. Pipeline 5 shows the least amount of variation across subjects; however, it shows together with Pipeline 3, the largest variation when studying the correlation between the nuisance norm and (**x** ∗ *w*)(**y** ∗ *w*) over all subjects and pairwise connections (Panel C) and **x** ∗ *w* over all subjects and regions (Panel D). Finally, it shows the lowest variation in (**x**^2^) ∗ *w* over all subjects and regions (Panel E). Fig. 9 shows equivalent violin plots using the time-varying correlation between the product of the windowed motion extracted from **x** and **y** obtained from Pipeline 3, instead of the nuisance norm. The results tell a similar story.

**Figure 9:**
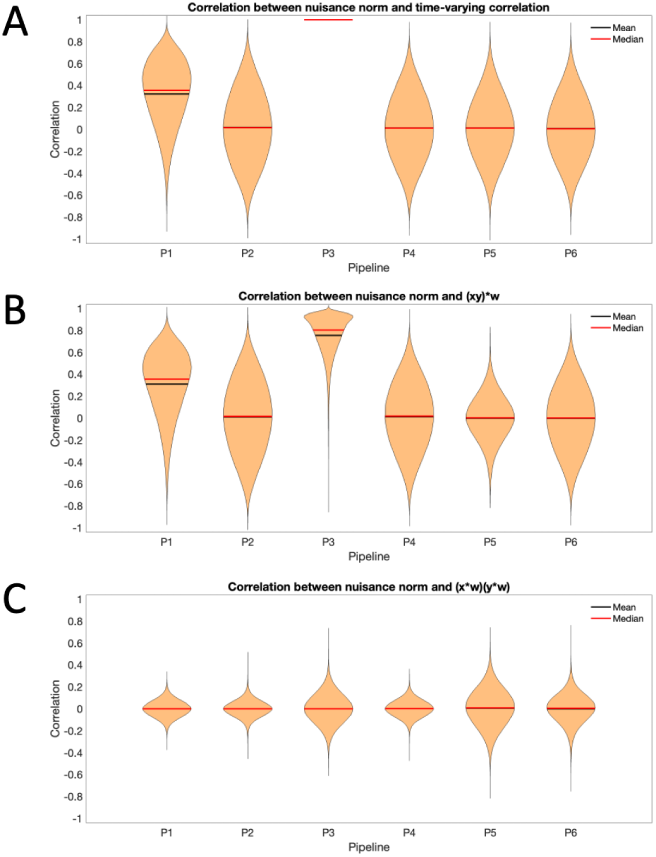
Correlation between the product of the windowed motion and the components of time-varying connectivity for Session 1. (A) The correlation between the product of the windowed motion and the time-varying connectivity plotted across all subjects and pair-wise connections. (B) The correlation between the product of the windowed motion and the time-varying value of (**xy**) ∗ *w* plotted across all subjects and pair-wise connections. (C) The correlation between the product of the windowed motion and the time-varying value of (**x** ∗ *w*)(**y** ∗ *w*) plotted across all subjects and pair-wise connections.

Fig. 10 shows the estimated brain states (left) and transition times (right) for each of the six pipelines. Starting with Pipeline 3 (third row), we see the states estimated using the nuisance images. The second state shows that most brain regions are highly interconnected, while the first state is significantly less connected. These states are further split into 8 separate functional networks as described in [16]: medial frontal (purple), frontoparietal (cyan), default mode (yellow), subcortical-cerebellum (red), motor (blue), visual I (pink), visual II (green), and visual association (maroon).

**Figure 10:**
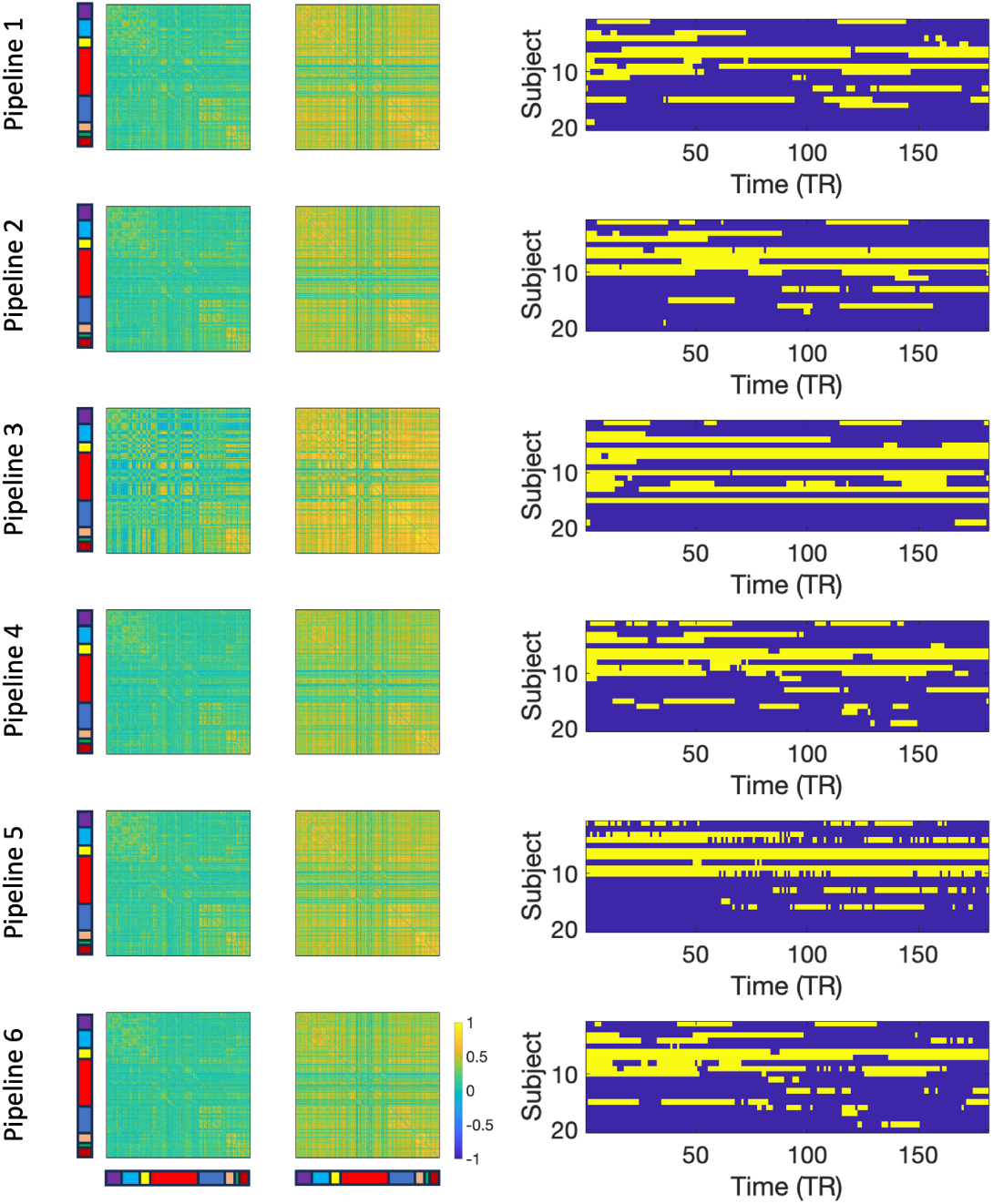
Results for Session 1. The two estimated brain states (left) and transition times between states (right) for each of the six pipelines. Each pipeline gives rise to two states, and the transitions between states across time are con-catenated across subjects giving a total of 180 × 21 time points.

While the data was concatenated across subjects prior to analysis, the analysis provided a measure of which state each subject was in at each time point. These results were rearranged into a subject-by-time matrix for easier viewing of transition times; see right panel.

Interestingly, the derived states from Pipeline 3 show a great deal of similarity to those found using the other Pipelines, which in many ways are not distinguishable from the nuisance states. The transition times for the three first pipelines are also similar, indicating that the transitions may be highly related to motion induced signal variation. The results from Pipeline 4-6 appear different as the correlations between regions are generally decreased and a number of cross-correlations between brain regions in State 2 are less apparent; perhaps changing the interpretation of the findings somewhat.

The similarity between Pipelines is further investigated in Fig. 11A. Here multidimensional scaling was applied to the twelve States; two states derived from each of the six Pipelines. The placement of the states in the first two dimensions is shown. Circles correspond to State 1 and squares to State 2, while the different pipelines are color-coded according to the figure legend. The green symbols correspond to the nuisance states (Pipeline 3). The States derived from Pipeline 1 (blue symbols) are close to them, while the States obtained after performing an initial motion correction (Pipeline 2; red symbols) lie yet further away. Finally, performing an additional motion regression moves the States yet further away from the nuisance states (Pipeline 4; cyan symbols), as does using a motion corrected filter (yellow symbols) and performing partial correlation (black symbols). Panel B shows a heatmap of the frame-wise displacement measured at every time point for every subject, illustrating the amount of motion for each subject. Finally, the left-hand side of Panel C shows the proportion nuisance present in the signal given by 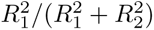. Here 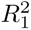 is the squared correlation between the motion data and the windowed time series and 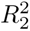 is the squared correlation between the motion-corrected data and the windowed time series. The results indicate a substantial amount of motion induced signal variation is present. The right-hand side of Panel C shows the same proportion averaged over each of the 8 functional networks. In particular, the subcortical-cerebellum and motor networks show significant impact of motion induced variation, with values ranging from 0.25 to 0.75.

**Figure 11:**
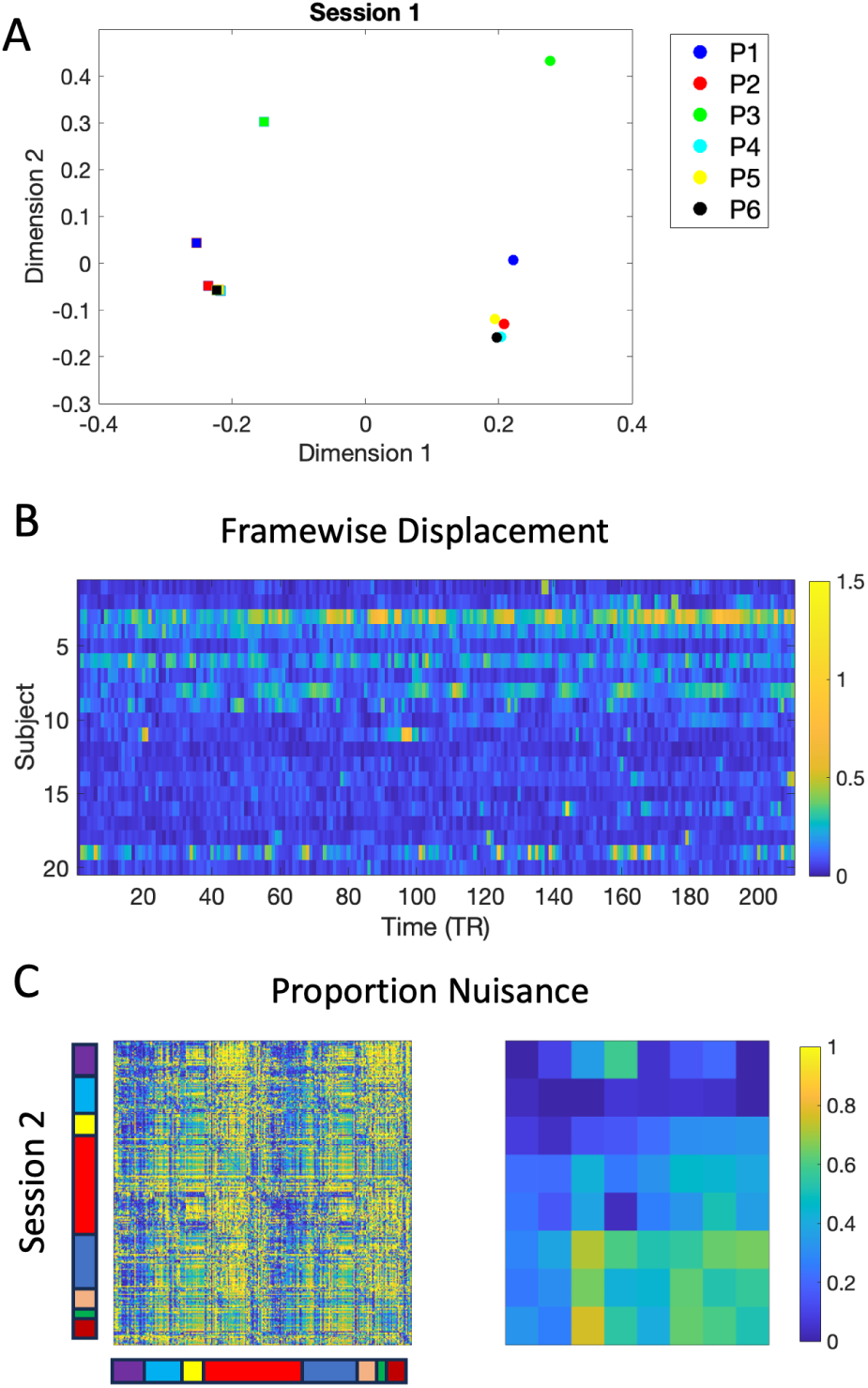
Results for Session 1. (A) Multidimensional scaling applied to the two states obtained from each of the six Pipelines. Circles correspond to State 1 and squares to State 2. The different pipelines are color-coded according to the figure legend. (B) A heatmap of the frame-wise displacement measured at every time point for every subject. (C) The proportion nuisance 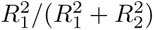, where 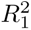 is the squared correlation between the motion data and the windowed time series and 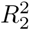 is the squared correlation between the motion-corrected data and the windowed time series. The left side shows the results for the 268 regions and the right side aggregated over the 8 networks.

Figure 12A-B shows the estimated distributions for the number of transitions and dwell time, respectively. Panel A shows the number of transitions, modeled using a Poisson distribution with a different *λ* value for each of the six pipelines, for Session 1. Referring back to the results seen in the multi-dimensional scaling plot (Fig. 11A), as states ‘move away’ from those observed in Pipeline 3, their distribution also begin to differ. The distributions for Pipelines 1 and 2 most resemble the distribution for Pipeline 3, indicating a smaller number of transitions, while those for Pipelines 4-6 show a right-ward shift indicating more transitions. Panel B shows the distribution for dwell time, which was modeled using an Exponential distribution with a different *µ* value for each of the six pipelines, for Session 1. This plot tells a similar story, which is not surprising as number of transitions and dwell time are highly related.

**Figure 12:**
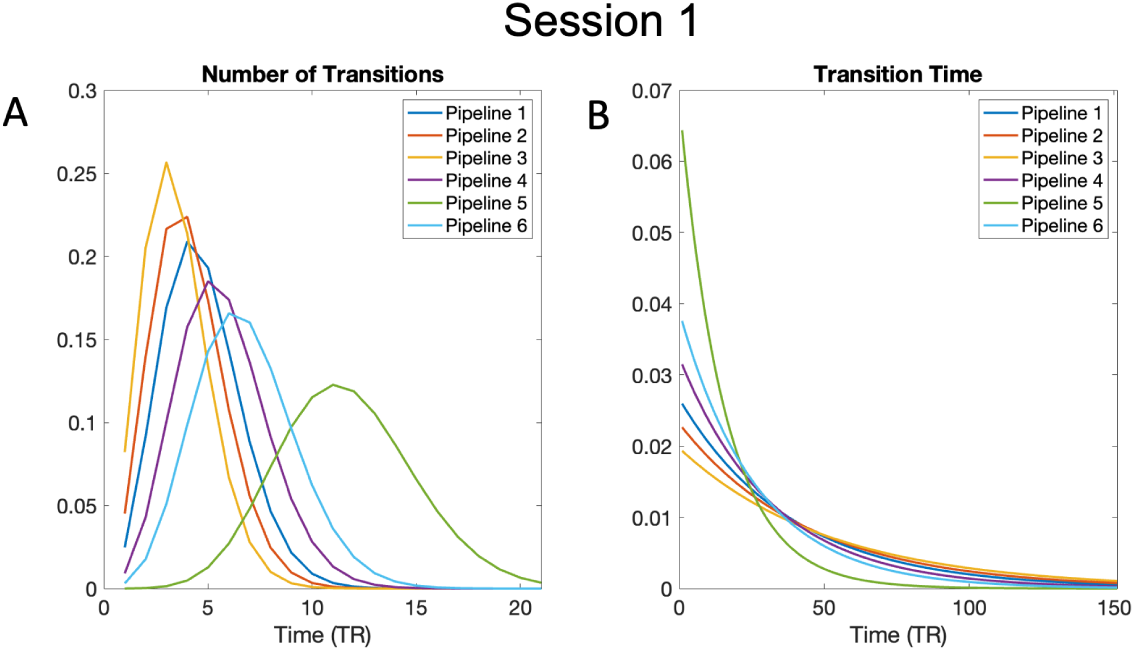
Estimated distributions for number of transitions and dwell time. (A) The number of subject-specific transitions modeled using a Poisson distribution for each of the six pipelines for Session 1. (B) The subject-specific dwell time, or the time spent in a state, modeled using an Exponential distribution for each of the six pipelines for Session 1.

Importantly, all of these results are reproducible using data obtained on the same subjects from a different scanning session. The equivalent results for Session 2 are seen in Figs. S1, S2, S3, S4, S5 and Fig. S6 which show strong similarity with the findings from the first session.

## 4 Discussion

In this paper we illustrate how sliding windows, a commonly used technique for assessing time-varying functional connectivity (TVFC) of rs-fMRI data, can interact with previous data preprocessing steps, thereby partially negating their effects. Sliding windows analysis entails convolving a signal in the time-domain with a rectangular (or tapered) filter. We show that this process may reintroduce correlation with motion induced signal variation, that was previously removed in the data cleaning portion of the analysis, and that these motion effects can potentially be mistaken for time-varying connectivity. Our analysis show that interactions between the application of the window and the nuisance regression and the fact that nuisance regression is not designed to be orthogonal to higher-order statistics both play important roles.

There has long been a debate over whether nuisance effects may be accounting for a significant portion of the observed fluctuations in TVFC. For example, [22] studied the effects of high-motion frames and came to the conclusion that much of the observed dynamics were related to motion. [20] proposed that nuisance effects may have larger impact when using a sliding-window approach due to the fact that transient effects can alter correlation estimates when using short windows. Finally, [27] showed that sliding windows correlation estimates can be strongly and significantly correlated with the sliding window norms associated with various nuisance measures. They further found that the correlation can persist even after nuisance regression was performed. They hypothesized several potential reasons for these observations, including the possibility that a large fraction of the nuisance term is orthogonal to the subspace spanned by the BOLD signals being studied. In this work we find equivalent results, but illustrate how it is caused by the interaction between the sliding window kernel and the nuisance regression. In addition, we show that standard nuisance regression approach may not be the optimal when studying time-varying higher-order statistics. To the best of our knowledge these links have not been noted in previous work.

This work complements previous work in the field ([18, 7, 24]) that showed how different preprocessing steps can interact with one another and thus counteract each other’s effect. The current work shows that perhaps more insidiously, later statistical analysis can similarly counteract previously performed preprocessing. We illustrate using a sliding window analysis, but the same problem should occur in any technique utilizing a moving window to assess TVFC. This includes alternative measures for assessing within-window connectivity such as partial correlations, coherence, and mutual information, as well as tapered sliding windows ([1]) and multivariate time series models such as the Dynamic Conditional Correlations ([25]).

This type of interaction is particularly problematic as the different components of data analysis have become increasingly compartmentalized. For example, in many research projects one group may acquire the data and do initial processing to place the data into a certain format, another group may preproccess or clean the data, while a third group may perform statistical analysis to the processed data. This compartmentalization has the effect of binning researchers into different non-overlapping silos, that may or may not actually communicate with one another even when working on the same dataset. This fragmentation makes it more difficult to safeguard against the issues raised in this paper.

How do we guard against such problems? We offer several suggestions. First, we find it helpful to run nuisance signal through the analysis pipeline and compare results with those obtained with the preprocessed data; see our Pipeline 3 for an example. This will allow researchers to study the similarity between findings to determine potential interactions.

Second, it is important to train researchers to understand the entire analytic pipeline from acquisition to final analysis. Ideally, the entire pipeline should be run together, with a careful examination of interactions between parts of the pipeline. If interactions are detected, these parts of the pipeline can be run simultaneously (e.g., as in Pipeline 4) or the different components can be orthogonalized with respect to one another. It is critical for researchers to critically evaluate commonly used analytic pipeline for flaws. We stress that these can sometimes be quite subtle and occur in unexpected ways.

Third, it should be possible to develop techniques for removing motion that do not interact with the filtering procedure. In this paper we investigate three alternative approaches for handling this issue. Two utilize block regression with varying numbers of regressors (Pipelines 4 and 6) and one uses a motion-corrected window function (Pipeline 5). Our investigations show that block regression generally outperforms the motion-corrected window function. The inclusion of additional regressors in the block regression in Pipeline 6 compared to Pipeline 4, does not appear to make a noticeable impact. However, while these approaches help mitigate the issue somewhat, none of them completely eliminate the problem. This is no doubt due to the fact that the projection performed within each window using truncated data and nuisance regressors, places the data in a subspace that need not be orthogonal to the subspace spanned by the full non-truncated data and nuisance regressors. In addition, the amount of data within a window makes performing motion regression in this manner difficult. In our illustration, we used a window length of 30 TR. If we perform motion regression consisting of 24 motion regressors this quickly uses all of the degrees of freedom available in the data. If we further seek to include additional covariates (e.g., CSF, white matter signal, global signal regression), we will have more regressors than observations. It may ultimately, be best to create an omnibus analysis technique that performs preprocessing and time-varying connectivity jointly. However, we leave the development of such a comprehensive approach to future work.

Our findings indicate that the correlation between the TVFC and nuisance signal is largely driven by the **xy** product. This is no doubt due to the fact that standard motion regression approaches are not designed for higher-order statistics. For example, the product of two motion-corrected time series will most likely be correlated with motion, as the Hadamard product need not be orthogonal to the space spanned by the motion regressors. This is seen in Fig, 2, and this correlation becomes increasingly important when studying time-varying measures of this metric. Ideally, one would use an approach that simultaneously performs motion correlation on all of the terms in Eq. 3, though the development of such an approach would be non-trivial.

A natural question to ask is whether one can avoid these problems by choosing an appropriate window length for the applied sliding-window. It may be possible to choose a window length that gives rise to a filter that minimally interacts with a certain type of nuisance signal. However, though we illustrated the problem using motion induced signal, we stress these issues can occur for any potential nuisance covariate, including scrubbing, component-based correction, physiological correction, and global signal regression. We anticipate that it would be difficult to safeguard against reintroduction of all of these artifacts by choosing a single ‘optimal’ window length.

How serious is this problem in practice? It is of course difficult to make a blanket statement on this issue. Fig. 11 provides an estimate of the proportion of nuisance-induced signal variation present in the signal. This metric allows us to quantify the proportion of motion induced signal variation present in the signal after the windowing procedure. Prior to windowing, this metric will be 0 in all regions of the brain. After windowing, the values are substantially higher. In particular, regions in the subcortical-cerebellum and motor networks show significant impact of motion induced signal variation, with values as high as 0.75 in Session 1 and 0.65 in Session 2.

Importantly, in our example data set, the nuisance data and the ’cleaned’ data give rise to similar state estimates and state transitions. This is concerning. It is naturally reasonable to question how ‘pure’ each of these signals are in practice. For example, the motion-corrected data presumably still includes some degree of motion induced signal, and similarly the nuisance images contain some neuronal signal. If these components cannot be adequately separated it may be the case that similarities in the state estimation are driven by this contamination. Regardless, we believe it is clear that the ultimate goal of preprocessing is to remove as much of this nuisance-induced signal variation as possible, after all otherwise why perform preprocessing in the first place? If so, it should not be considered acceptable to apply a statistical technique that potentially negates this effort. As a field we should seek to develop better methods that don’t allow for this possibility.

Ultimately, a recommendation is to avoid using window-based measures of TVFC, and instead focus on using measures that directly estimate brain states such as change point analysis ([12, 13, 37]) or variants of hidden Markov models (HMMs) ([15, 36, 31, 6]). At the very least, researchers should complement their window-based approaches with these types of analysis, as well as repeat their analyses using nuisance signal (as done in Pipeline 3). Together, this will allow for more careful deliberation whether results are spurious or not.

In summary, this work highlights the potential for correlations with confounding variables being reintroduced into the data when sliding windows analysis is performed. This may negate the assumption that these effects are adequately controlled in preprocessing. The results raise the question to what degree estimated brain states reflect non-neuronal nuisance signal variation (e.g., related to motion) rather than the neuronal signal of interest.

## Data Availability statement

All code used in this paper is available at https://github.com/mal2053/SlidingWindowMotion/. The data used in this paper is available by contacting the author.

## Declarations of competing Interests

The author declares he has no known competing interests.

## Contributions

MAL performed all work on this manuscript.

## Acknowledgments

The work presented in this paper was supported in part by NIH grants R01 EB026549 from the National Institute of Biomedical Imaging and Bioengineering and R01 MH129397 from the National Institute of Mental Health. The author thanks Brian Caffo, Ciprian Crainiceanu, Robert Welsh, and Jim Pekar for helpful comments on the manuscript.

## Supplementary Material

**Figure S1:**
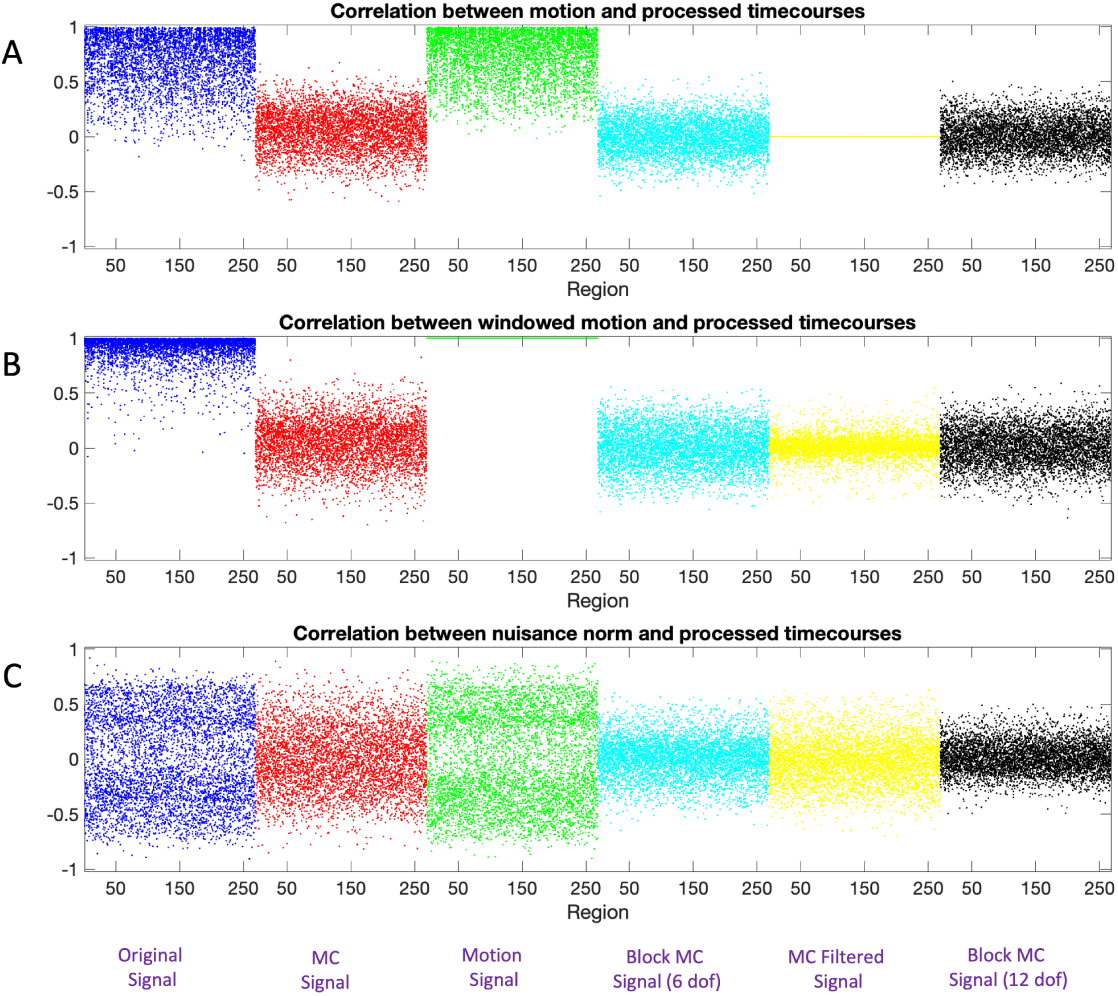
Correlation between motion and windowed rs-fMRI signal for Session 2. (A) The correlation between the motion signal and the windowed rs-fMRI signal for each of the 20 subjects shown for each region and pipeline. Note there are 20 dots representing the subject-specific values at each point on the x-axis. (B) The correlation between the windowed motion signal and the windowed rs-fMRI signal for each of the 20 subjects shown for each region and pipeline. (C) The correlation between the nuisance norm and the windowed rs-fMRI signal for each of the 20 subjects shown for each region and pipeline. The color-coded as follows: Pipeline 1 (Blue), Pipeline 2 (Red), Pipeline 3 (Green), Pipeline 4 (Cyan), and Pipeline 5 (Yellow).

**Figure S2:**
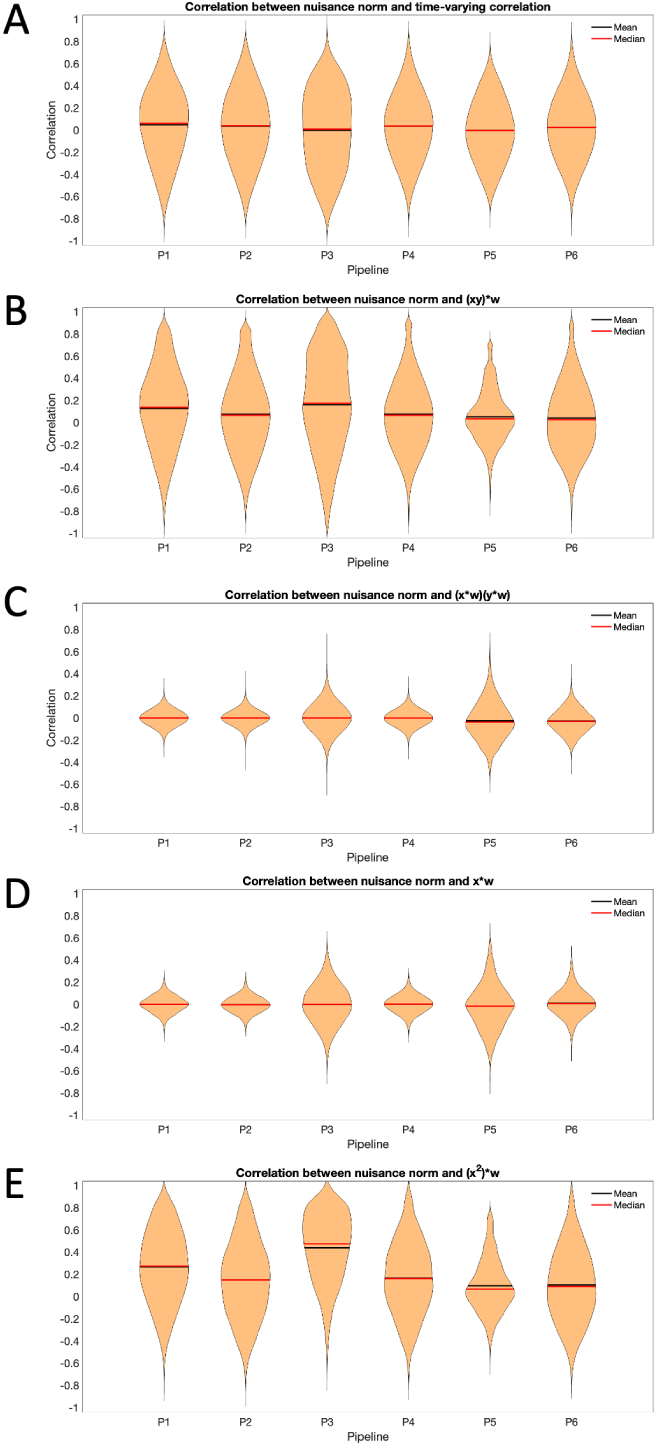
Correlation between the nuisance norm and the components of time-varying connectivity for Session 2. (A) The correlation between the nuisance norm and the time-varying connectivity plotted across all subjects and pair-wise connections. (B) The correlation between the nuisance norm and the time-varying value of (**xy**) ∗ *w*, where **x** and **y** are the signals of interest, plotted across all subjects and pair-wise connections. (C) The correlation between the nuisance norm and the time-varying value of (**x** ∗ *w*)(**y** ∗ *w*) plotted across all subjects and pair-wise connections. (D) The correlation between the nuisance norm and the time-varying value of (**x** ∗ *w*) plotted across all subjects and regions. (E) The correlation between the nuisance norm and the time-varying value of (**x**^2^) ∗ *w* plotted across all subjects and regions.

**Figure S3:**
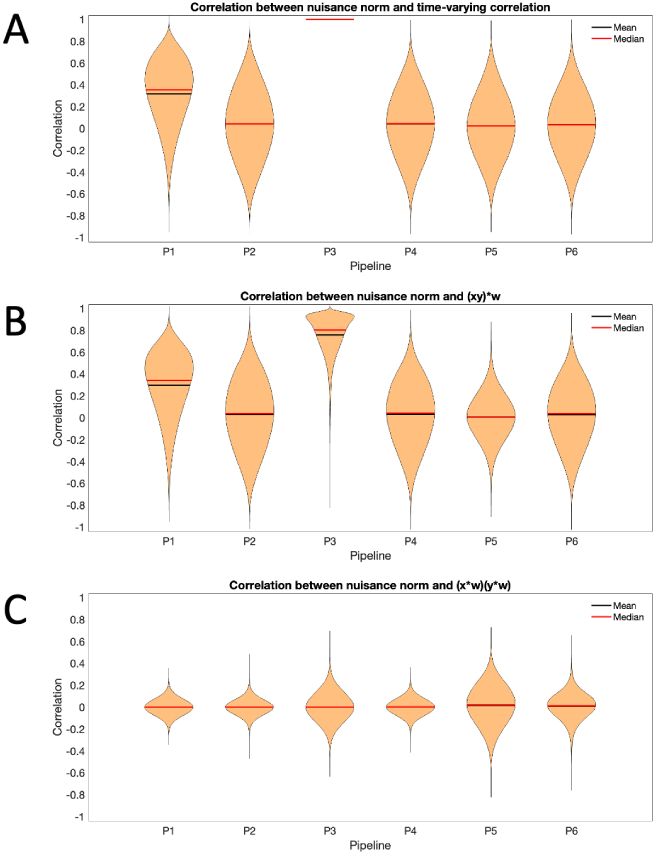
Correlation between the product of the windowed motion and the components of time-varying connectivity for Session 2. (A) The correlation between the product of the windowed motion and the time-varying connectivity plotted across all subjects and pair-wise connections. (B) The correlation between the product of the windowed motion and the time-varying value of (**xy**) ∗ *w* plotted across all subjects and pair-wise connections. (C) The correlation between the product of the windowed motion and the time-varying value of (**x** ∗ *w*)(**y** ∗ *w*) plotted across all subjects and pair-wise connections.

**Figure S4:**
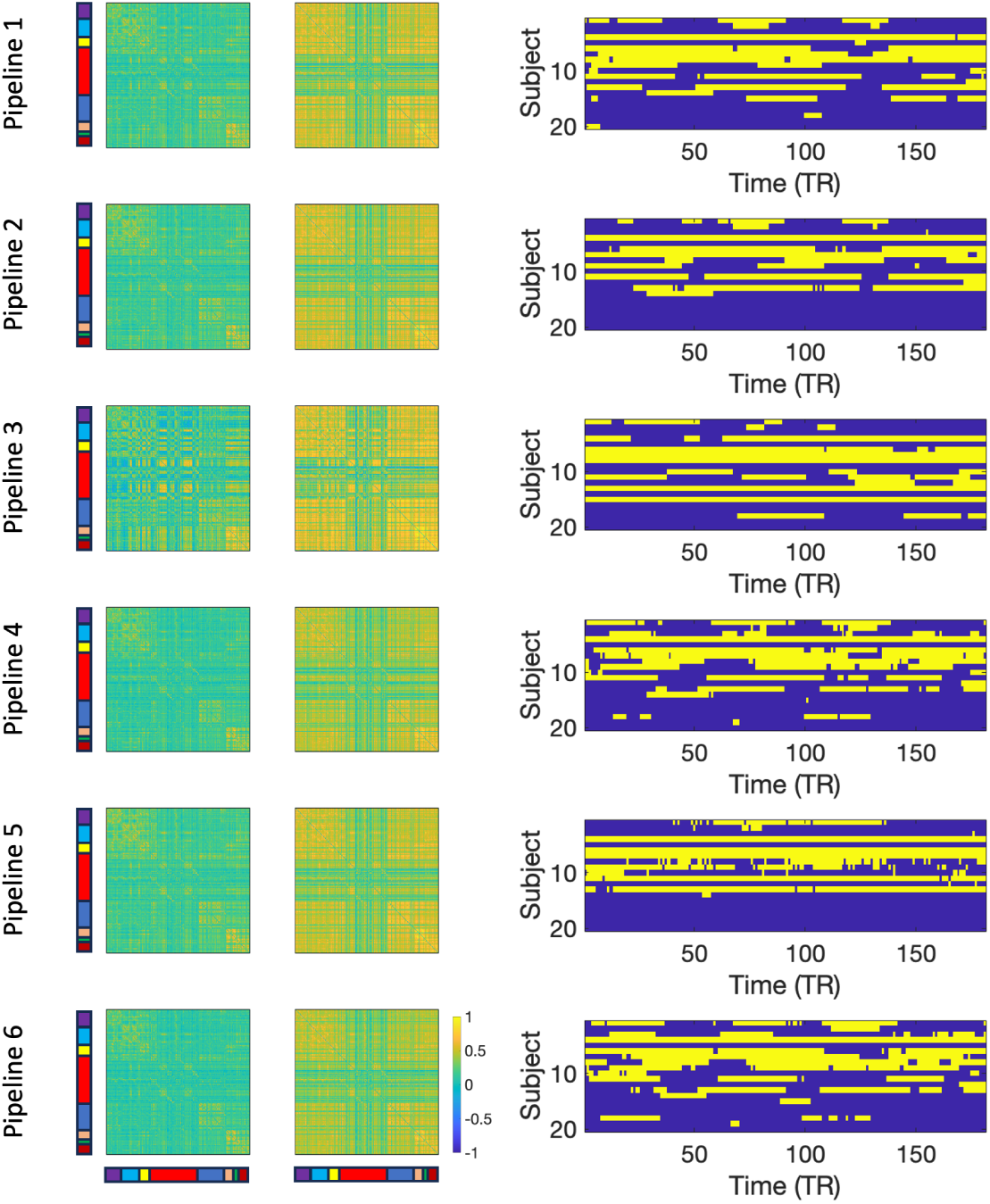
Results for Session 2. The two estimated brain states (left) and transition times between states (right) for each of the four pipelines. Each pipeline gives rise to two states, and the transitions between states across time are concatenated across subjects giving a total of 180 × 21 time points.

**Figure S5:**
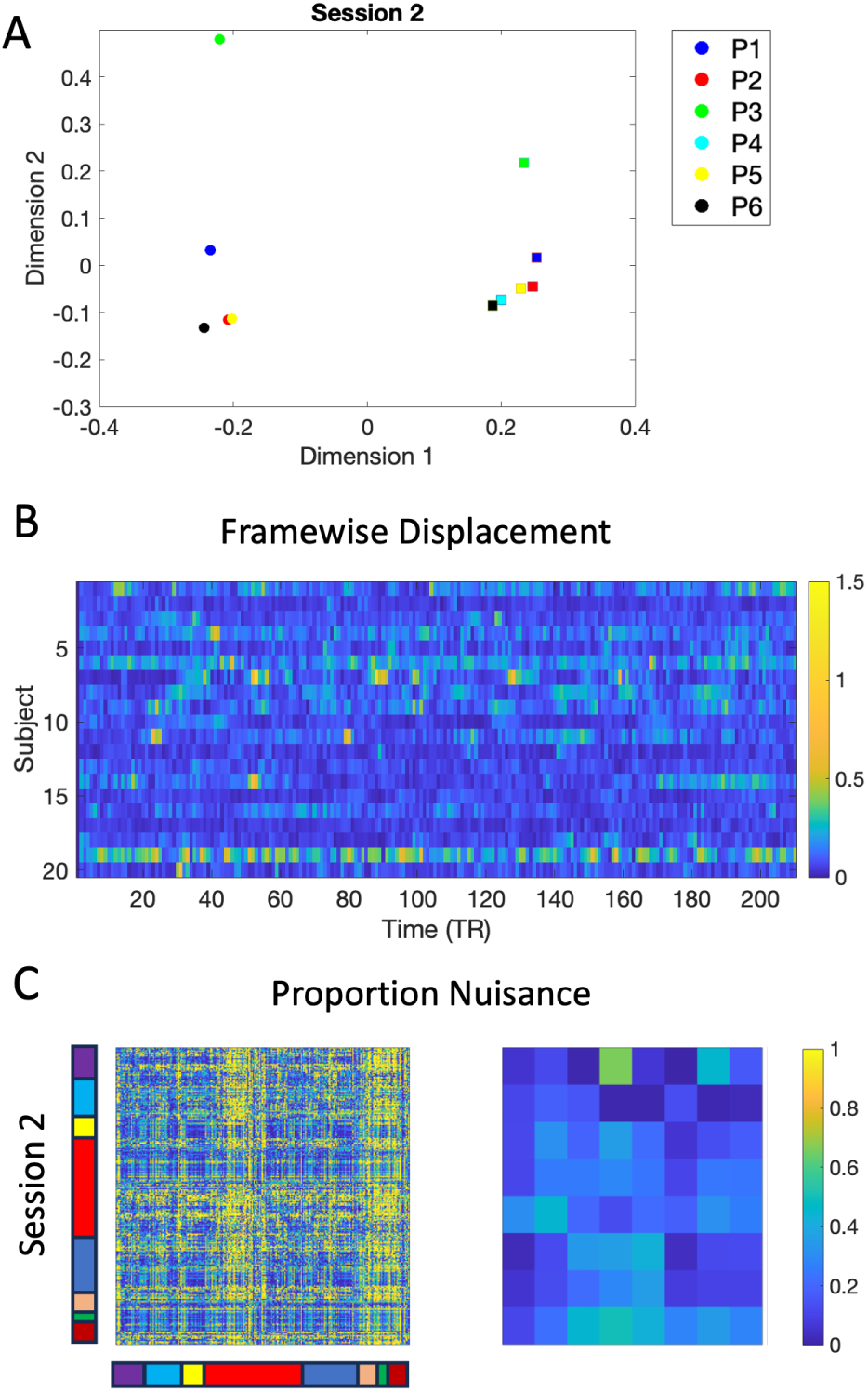
Results for Session 2. (A) Multidimensional scaling applied to the two states obtained from each of the six Pipelines. Circles correspond to State 1 and squares to State 2. The different pipelines are color-coded according to the figure legend. (B) A heatmap of the frame-wise displacement measured at every time point for every subject. (C) The proportion nuisance 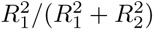, where 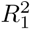 is the squared correlation between the motion data and the windowed time series and 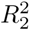 is the squared correlation between the motion-corrected data and the windowed time series. The left side shows the results for the 268 regions and the right side aggregated over the 8 networks.

**Figure S6:**
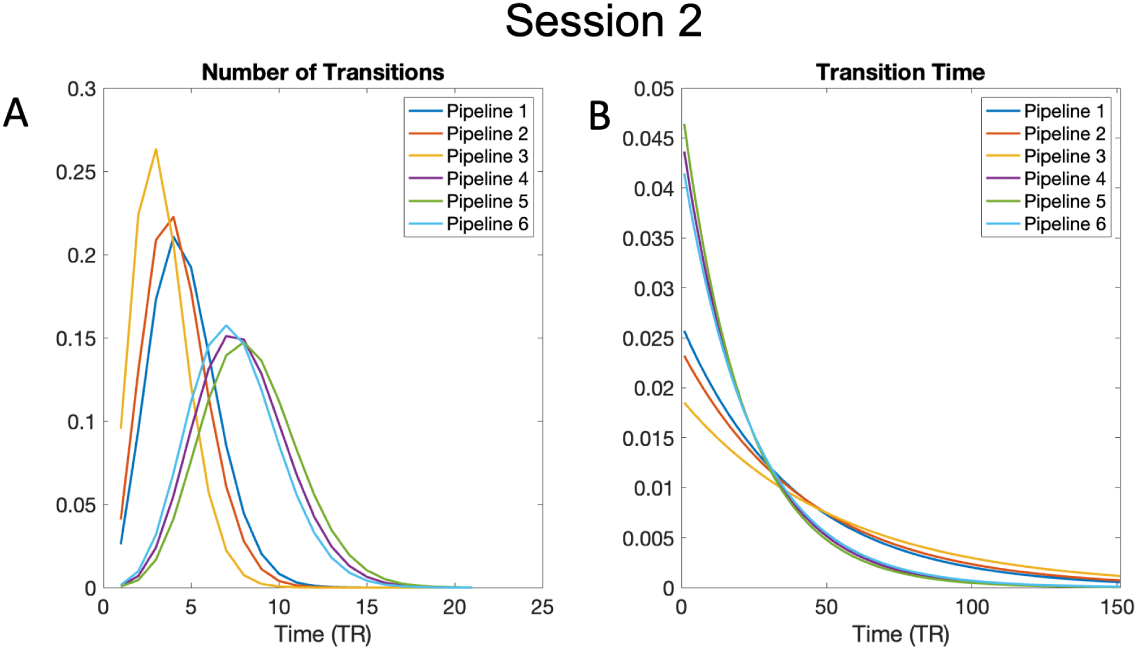
Estimated distributions for number of transitions and dwell time. (A) The number of subject-specific transitions modeled using a Poisson distribution for each of the six pipelines for Session 2. (B) The subject-specific dwell time, or the time spent in a state, modeled using an Exponential distribution for each of the six pipelines for Session 2.

## Footnotes

1 Publicly available at http://www.nitrc.org/projects/multimodal

## Notes

### Competing Interest Statement

The authors have declared no competing interest.

### Summary of Updates

This version of the manuscript has been updated to clarify a number of theoretical issues, and further explore the relationship between various components of the time-varying correlation with motion. In addition, a simulation study has been added.

## References

[1] E. A. Allen, E. Damaraju, S. M. Plis, E. B. Erhardt, T. Eichele, and V. D. Calhoun. Tracking whole-brain connectivity dynamics in the resting state. Cerebral cortex, 24(3):663–676, 2014.

[2] J. Ashburner and K. J. Friston. Unified segmentation. Neuroimage, 26(3):839–851, 2005.

[3] C. F. Beckmann, M. DeLuca, J. T. Devlin, and S. M. Smith. Investigations into resting-state connectivity using independent component analysis. Philosophical Transactions of the Royal Society B: Biological Sciences, 360(1457):1001–1013, 2005.

[4] R. M. Birn, J. B. Diamond, M. A. Smith, and P. A. Bandettini. Separating respiratory-variation-related fluctuations from neuronal-activity-related fluctuations in fmri. Neuroimage, 31(4):1536–1548, 2006.

[5] B. Biswal, F. Zerrin Yetkin, V. M. Haughton, and J. S. Hyde. Functional connectivity in the motor cortex of resting human brain using echo-planar mri. Magnetic resonance in medicine, 34(4):537–541, 1995.

[6] T. A. Bolton, A. Tarun, V. Sterpenich, S. Schwartz, and D. Van De Ville. Interactions between large-scale functional brain networks are captured by sparse coupled hmms. IEEE transactions on medical imaging, 37(1):230–240, 2018.

[7] M. G. Bright, C. R. Tench, and K. Murphy. Potential pitfalls when denoising resting state fmri data using nuisance regression. Neuroimage, 154:159–168, 2017.

[8] C. Caballero-Gaudes and R. C. Reynolds. Methods for cleaning the bold fmri signal. Neuroimage, 154:128–149, 2017.

[9] V. D. Calhoun, R. Miller, G. Pearlson, and T. Adalı. The chronnectome: time-varying connectivity networks as the next frontier in fmri data discovery. Neuron, 84(2):262–274, 2014.

[10] C. Chang and G. H. Glover. Time–frequency dynamics of resting-state brain connectivity measured with fmri. Neuroimage, 50(1):81–98, 2010.

[11] R. Ciric, D. H. Wolf, J. D. Power, D. R. Roalf, G. L. Baum, K. Ruparel, R. T. Shinohara, M. A. Elliott, S. B. Eickhoff, C. Davatzikos, et al. Bench-marking of participant-level confound regression strategies for the control of motion artifact in studies of functional connectivity. Neuroimage, 154:174–187, 2017.

[12] I. Cribben, R. Haraldsdottir, L. Y. Atlas, T. D. Wager, and M. A. Lindquist. Dynamic connectivity regression: determining state-related changes in brain connectivity. Neuroimage, 61(4):907–920, 2012.

[13] I. Cribben, T. D. Wager, and M. A. Lindquist. Detecting functional connectivity change points for single-subject fmri data. Frontiers in Computational Neuroscience, 7, 2013.

[14] M. De Luca, C. Beckmann, N. De Stefano, P. Matthews, and S. M. Smith. fmri resting state networks define distinct modes of long-distance interactions in the human brain. Neuroimage, 29(4):1359–1367, 2006.

[15] H. Eavani, T. D. Satterthwaite, R. E. Gur, R. C. Gur, and C. Davatzikos. Unsupervised learning of functional network dynamics in resting state fmri. In International Conference on Information Processing in Medical Imaging, pages 426–437. Springer, 2013.

[16] E. S. Finn, X. Shen, D. Scheinost, M. D. Rosenberg, J. Huang, M. M. Chun, X. Papademetris, and R. T. Constable. Functional connectome fingerprinting: identifying individuals using patterns of brain connectivity. Nature neuroscience, 18(11):1664, 2015.

[17] K. J. Friston, S. Williams, R. Howard, R. S. Frackowiak, and R. Turner. Movement-related effects in fmri time-series. Magnetic resonance in medicine, 35(3):346–355, 1996.

[18] M. N. Hallquist, K. Hwang, and B. Luna. The nuisance of nuisance regression: spectral misspecification in a common approach to resting-state fmri preprocessing reintroduces noise and obscures functional connectivity. Neuroimage, 82:208–225, 2013.

[19] D. A. Handwerker, V. Roopchansingh, J. Gonzalez-Castillo, and P. A. Bandettini. Periodic changes in fmri connectivity. Neuroimage, 63(3):1712–1719, 2012.

[20] R. M. Hutchison, T. Womelsdorf, E. A. Allen, P. A. Bandettini, V. D. Calhoun, M. Corbetta, S. Della Penna, J. H. Duyn, G. H. Glover, J. Gonzalez-Castillo, et al. Dynamic functional connectivity: promise, issues, and interpretations. Neuroimage, 80:360–378, 2013.

[21] B. A. Landman, A. J. Huang, A. Gifford, D. S. Vikram, I. A. L. Lim, J. A. Farrell, J. A. Bogovic, J. Hua, M. Chen, S. Jarso, et al. Multiparametric neuroimaging reproducibility: a 3-t resource study. Neuroimage, 54(4):2854–2866, 2011.

[22] T. O. Laumann, A. Z. Snyder, A. Mitra, E. M. Gordon, C. Gratton, B. Adeyemo, A. W. Gilmore, S. M. Nelson, J. J. Berg, D. J. Greene, et al. On the stability of bold fmri correlations. Cerebral cortex, 27(10):4719–4732, 2017.

[23] N. Leonardi and D. Van De Ville. On spurious and real fluctuations of dynamic functional connectivity during rest. Neuroimage, 104:430–436, 2015.

[24] M. A. Lindquist, S. Geuter, T. D. Wager, and B. S. Caffo. Modular preprocessing pipelines can reintroduce artifacts into fmri data. Human brain mapping, 40(8):2358–2376, 2019.

[25] M. A. Lindquist, Y. Xu, M. B. Nebel, and B. S. Caffo. Evaluating dynamic bivariate correlations in resting-state fmri: a comparison study and a new approach. NeuroImage, 101:531–546, 2014.

[26] D. J. Lurie, D. Kessler, D. S. Bassett, R. F. Betzel, M. Breakspear, S. Kheilholz, A. Kucyi, R. Liégeois, M. A. Lindquist, A. R. McIntosh, et al. Questions and controversies in the study of time-varying functional connectivity in resting fmri. Network neuroscience, 4(1):30–69, 2020.

[27] A. Nalci, B. D. Rao, and T. T. Liu. Nuisance effects and the limitations of nuisance regression in dynamic functional connectivity fmri. Neuroimage, 184:1005–1031, 2019.

[28] J. D. Power, A. Mitra, T. O. Laumann, A. Z. Snyder, B. L. Schlaggar, and S. E. Petersen. Methods to detect, characterize, and remove motion artifact in resting state fmri. Neuroimage, 84:320–341, 2014.

[29] M. G. Preti, T. A. Bolton, and D. Van De Ville. The dynamic functional connectome: State-of-the-art and perspectives. Neuroimage, 160:41–54, 2017.

[30] K. P. Pruessmann, M. Weiger, M. B. Scheidegger, P. Boesiger, et al. Sense: sensitivity encoding for fast mri. Magnetic resonance in medicine, 42(5):952–962, 1999.

[31] H. M. Shappell, B. S. Caffo, J. J. Pekar, and M. Lindquist. Improved state change estimation in dynamic functional connectivity using hidden semi-markov models. bioRxiv, page 519868, 2019.

[32] X. Shen, F. Tokoglu, X. Papademetris, and R. T. Constable. Groupwise whole-brain parcellation from resting-state fmri data for network node identification. Neuroimage, 82:403–415, 2013.

[33] K. Shmueli, P. van Gelderen, J. A. de Zwart, S. G. Horovitz, M. Fukunaga, J. M. Jansma, and J. H. Duyn. Low-frequency fluctuations in the cardiac rate as a source of variance in the resting-state fmri bold signal. Neuroimage, 38(2):306–320, 2007.

[34] M. K. Stehling, R. Turner, and P. Mansfield. Echo-planar imaging: magnetic resonance imaging in a fraction of a second. Science, 254(5028):43–50, 1991.

[35] K. R. Van Dijk, M. R. Sabuncu, and R. L. Buckner. The influence of head motion on intrinsic functional connectivity mri. Neuroimage, 59(1):431–438, 2012.

[36] D. Vidaurre, S. M. Smith, and M. W. Woolrich. Brain network dynamics are hierarchically organized in time. Proceedings of the National Academy of Sciences, 114(48):12827–12832, 2017.

[37] Y. Xu and M. A. Lindquist. Dynamic connectivity detection: an algorithm for determining functional connectivity change points in fmri data. Frontiers in Neuroscience, 9, 2015.

[38] C. Yan and Y. Zang. Dparsf: a matlab toolbox for” pipeline” data analysis of resting-state fmri. Frontiers in systems neuroscience, 4:13, 2010.

